# *Sinorhizobium meliloti* functions required for resistance to antimicrobial NCR peptides and bacteroid differentiation

**DOI:** 10.1101/2020.12.04.412775

**Authors:** Quentin Nicoud, Quentin Barrière, Nicolas Busset, Sara Dendene, Dmitrii Travin, Mickaël Bourge, Romain Le Bars, Claire Boulogne, Marie Lecroël, Sándor Jenei, Atilla Kereszt, Eva Kondorosi, Emanuele G. Biondi, Tatiana Timchenko, Benoît Alunni, Peter Mergaert

**Affiliations:** Université Paris-Saclay, CEA, CNRS, Institute for Integrative Biology of the Cell (I2BC), Gif-sur-Yvette, France; Center of Life Sciences, Skolkovo Institute of Science and Technology, Moscow, Russia; Institute of Plant Biology, Biological Research Centre, Szeged, Hungary

**Author notes:** equal contribution (Quentin Nicoud, Quentin Barrière, Nicolas Busset).

## Abstract

Legumes of the *Medicago* genus form symbiosis with the bacterium *Sinorhizobium meliloti* and develop root nodules housing large numbers of the intracellular symbionts. Members of the Nodule-specific Cysteine Rich peptide (NCRs) family induce the endosymbionts into a terminal differentiated state. Individual cationic NCRs are antimicrobial peptides that have the capacity to kill the symbiont but the nodule cell environment prevents killing. Moreover, the bacterial broad-specificity peptide uptake transporter BacA and exopolysaccharides contribute to protect the endosymbionts against the toxic activity of NCRs. Here, we show that other *S. meliloti* functions participate in the protection of the endosymbionts, including an additional broad-specificity peptide uptake transporter encoded by the *yejABEF* genes, lipopolysaccharide modifications mediated by *lpsB* and *lpxXL* as well as *rpoH1*, encoding a stress sigma factor. Mutants of these genes show *in vitro* a strain-specific increased sensitivity profile against a panel of NCRs and form nodules in which bacteroid differentiation is affected. The *lpsB* mutant nodule bacteria do not differentiate, the *lpxXL* and *rpoH1* mutants form some seemingly fully differentiated bacteroids although most of the nodule bacteria are undifferentiated, while the *yejABEF* mutants form hypertrophied but nitrogen-fixing bacteroids. The nodule bacteria of all the mutants have a strongly enhanced membrane permeability, which is dependent on the transport of NCRs to the endosymbionts. Our results suggest that *S. meliloti* relies on a suite of functions including peptide transporters, the bacterial envelope structures and stress response regulators to resist the aggressive assault of NCR peptides in the nodule cells.

**Importance:** The nitrogen fixing symbiosis of legumes with rhizobium bacteria has a predominant ecological role in the nitrogen cycle and has the potential to provide the nitrogen required for plant growth in agriculture. The host plants allow the rhizobia to colonize specific symbiotic organs, the nodules, in large numbers in order to produce sufficient reduced nitrogen for the plant needs. Some legumes, including *Medicago* spp., produce massively antimicrobial peptides to keep this large bacterial population in check. These peptides, known as NCRs, have the potential to kill the rhizobia but in nodules, they rather inhibit the division of the bacteria, which maintain a high nitrogen fixing activity. In this study, we show that the tempering of the antimicrobial activity of the NCR peptides in the *Medicago* symbiont *Sinorhizobium meliloti* is multifactorial and requires the YejABEF peptide transporter, the lipopolysaccharide outer membrane composition and the stress response regulator RpoH1.

## Introduction

Antimicrobial peptides (AMPs) are essential mediators of innate immunity in eukaryotes. Their function is to attack and kill harmful invading microbes. Animals deficient in a specific AMP or in the regulation of AMP expression are more sensitive to bacterial or fungal infections, while the ectopic expression of AMPs in animals and plants enhances resistance (1–3). Many organisms have also recruited AMPs as essential regulators of bacteria in symbiotic associations (2). In symbiosis, hosts intentionally maintain bacterial partners and the role of “symbiotic” AMPs is therefore not to eradicate the symbiotic microbes but rather to police them or to optimize their metabolic integration with the hosts (2, 4).

An extreme case of deployment of AMPs for controlling endosymbiont populations, involving hundreds of peptides, is described in the rhizobium-legume symbiosis (2, 5–9). Legumes form a symbiosis with phylogenetically diverse nitrogen-fixing soil bacteria, collectively called rhizobia. This nutritional symbiosis provides reduced nitrogen to the plants, enabling them to grow in nitrogen-poor soils. The symbiosis implies the formation of nodules, specific symbiotic organs, on the roots of the plants. These nodules house the nitrogen-fixing rhizobia, which transfer their produced ammonia to the plant in return for the exclusive niche in the nodules where they multiply massively from a single or very few infecting bacteria to a population of millions.

After endocytic uptake by the symbiotic nodule cells, the multiplied bacteria reside intracellularly in vesicles called symbiosomes. The nodule cells and symbiosomes establish the optimal conditions for nitrogen fixation and metabolic exchange with the endosymbionts and at the same time, keep them in check. The low oxygen levels prevailing in the symbiotic nodule cells transform the rhizobia into a differentiated physiological state, called bacteroid, which is adapted for nitrogen fixation. Moreover, in certain legume clades like the Inverted Repeat Lacking Clade (IRLC) and the Dalbergioids, the physiological transition of the bacteroids is accompanied with a remarkable differentiation process that is manifested in an irreversible loss of the capacity of bacteroids to divide (10, 11). These terminally differentiated bacteroids have a partially permeabilized cell membrane. They are giant bacterial cells. A switch in the bacterial cell cycle, from a regular succession of replication and division to a series of repeated genome replications without divisions, drives this cell enlargement, resulting in polyploid bacteroids.

The terminal differentiation is triggered by a family of Nodule-specific Cysteine-Rich peptides (NCRs), produced by the symbiotic nodule cells (11–13). Over 600 *NCR* genes were identified in the *Medicago truncatula* genome, while the NCR repertoire in other species of the IRLC and the Dalbergoids ranges from a few to several hundred (11, 12, 14). The *NCR* genes in *M. truncatula* are specifically expressed in the symbiotic cells (15). They are activated in waves during the differentiation of the bacteroids, including sets of *NCR* genes activated at the onset and others at the intermediate or final stages of the differentiation. These temporal profiles indicate that the *NCR* genes have specific functions during the bacteroid formation process.

The NCR peptides have structural features shared with AMPs and at least some NCRs, in particular the cationic ones, can kill or inhibit *in vitro* the growth of not only the rhizobium symbionts but also many other bacteria and even fungi (16). Their major antibacterial activity results from their capacity to disturb the integrity of the inner and outer membranes of bacteria (13, 17), although some NCRs also have inhibiting activities on essential intracellular machineries (18). However, bacteroids remain metabolically active for a very long time, despite the high burden of NCRs. Possibly, the environment of the symbiotic nodule cells and symbiosomes contributes to tempering the antimicrobial activity of the peptides. Importantly, also specific functions of the bacteria themselves are NCR-resistance determinants in the bacteroids.

*Sinorhizobium meliloti*, the symbiont of *Medicago* plants, requires the peptide transporter BacA to counter the NCR peptides inside the symbiotic nodule cells and establish a chronic infection (19). *S. meliloti bacA* mutants are hypersensitive to the antimicrobial NCRs. They induce nodules and infect plant cells seemingly normally but the mutants die rapidly after their release in the symbiotic cells. This death can be avoided by blocking NCR transport to the infecting rhizobia in the *M. truncatula dnf1* mutant (19). BacA proteins are broad-specificity peptide uptake transporters (20–22). They can promote the uptake of NCR peptides, suggesting that BacA provides resistance by redirecting the peptides away from the bacterial membrane, thereby limiting membrane damage. Exopolysaccharide (EPS) is another known factor of *S. meliloti* that helps the endosymbionts to withstand the NCRs (14, 23). This negatively charged extracellular polysaccharide traps the cationic AMPs, reducing their effective concentration in the membrane vicinity.

Bacterial resistance to AMPs is usually multifactorial (24), suggesting that besides BacA and EPS, additional functions of *S. meliloti* bacteroids contribute to resisting the NCRs in the symbiotic nodule cells. The literature on *S. meliloti* is rich in the description of bacterial genes that are required for symbiosis. However, the reporting on these mutants often lacks precise information on their bacteroid phenotype and/or on their sensitivity to NCRs. Moreover, transcriptome and Tn-seq (transposon sequencing) analyses of NCR-treated cells and NCR-protein interaction studies identified a whole suite of additional candidate NCR-responsive functions in *S. meliloti* (18, 25–27). Together with BacA and EPS, some of these *S. meliloti* functions may contribute to alleviate the NCR stress on the bacteroids. To test this hypothesis, we have selected in the present study three candidate functions and analyzed the phenotype of the corresponding mutants in NCR resistance and bacteroid formation.

## Results

### *Sinorhizobium meliloti* mutants with enhanced sensitivity to NCR peptides

The *S. meliloti* functions selected in this study include a broad-specificity peptide uptake transporter encoded by the *yejABEF* genes (SMc02829-SMc02832), lipopolysaccharide (LPS) modifications mediated by *lpsB* (SMc01219) and *lpxXL* (SMc04268) as well as *rpoH1* (SMc00646), encoding a stress sigma factor. The YejABEF ABC transporter was selected on the basis of a genetic screen by Tn-seq in *S. meliloti* revealing that mutants have an increased sensitivity to the peptide NCR247 (25). LPS structure is one of the major determinants of AMP resistance and sensitivity in Gram-negative bacteria (24, 28). The selected genes *lpsB* and *lpxXL* encode a glycosyltransferase involved in the synthesis of the LPS core and a very-long-chain fatty acid acyltransferase involved in the biosynthesis of lipid A, respectively. Mutants in these genes are affected in resistance to AMPs and in symbiosis (29, 30). Finally, the *rpoH1* gene is a global stress response regulator in *S. meliloti* (31). This gene as well as its target genes are upregulated in NCR247 treated cells (26, 27). An *rpoH1* mutant is also affected in symbiosis (32). These selected genes are expressed in nodules, with peak expression in different regions of the nodule where bacteria infect plant cells, undergo the differentiation process or fix nitrogen (15, 33)(Fig. S1).

The sensitivity of mutants in these candidate genes was tested with a small panel of cationic NCR peptides that were previously shown to have antimicrobial activity (14). The tested peptides, NCR169 (isoelectric point 8.45), NCR183 (10.10), NCR247 (10.15) and NCR280 (9.80), displayed three different expression patterns in the nodule tissues (15). The *NCR280* gene is expressed in the younger nodule cells, the *NCR169* gene in the older cells while *NCR183* and *NCR247* have an intermediate expression pattern (Fig. S1). The selected mutants were tested along with the wild-type strain and the *bacA* mutant, which was previously shown to be hypersensitive to NCRs (19). The four tested NCR peptides had a strong antimicrobial activity against the wild-type strain, which displayed a survival rate ranging from 8% to 0.03% depending on the tested peptide (Table S1). In agreement with previous results, the *bacA* mutant was hypersensitive to the four peptides (Table S1; Fig. 1). Interestingly, the newly analyzed mutants all displayed a higher sensitivity to at least one of the peptides compared to the wild type (Table S1; Fig. 1). The *lpxXL* mutant was more sensitive to the four peptides, the *lpsB, yejE* and *yejF* mutants were more sensitive to NCR183, NCR247 and NCR280. The *yejA* mutant was more sensitive to NCR280 and the *rpoH1* mutant was more sensitive to NCR247. The differential response towards peptide NCR247 of the *yejA* mutant on the one hand and the *yejE* and *yejF* mutants on the other hand, corresponds to the previously described Tn-seq screen with this peptide (25). Contrary to the other mutants, the mutants in the YejABEF transporter genes were newly constructed and not characterized before. Therefore, we confirmed that the NCR247 sensitivity phenotype was directly attributable to the inactivation of the transporter by complementation of the *yejF* mutant phenotype with a plasmid-borne copy of the *yejABEF* genes (Table S1). Taken together, our analysis indicated that each mutant displayed a specific sensitivity profile to the panel of tested peptides.

**Figure 1.**
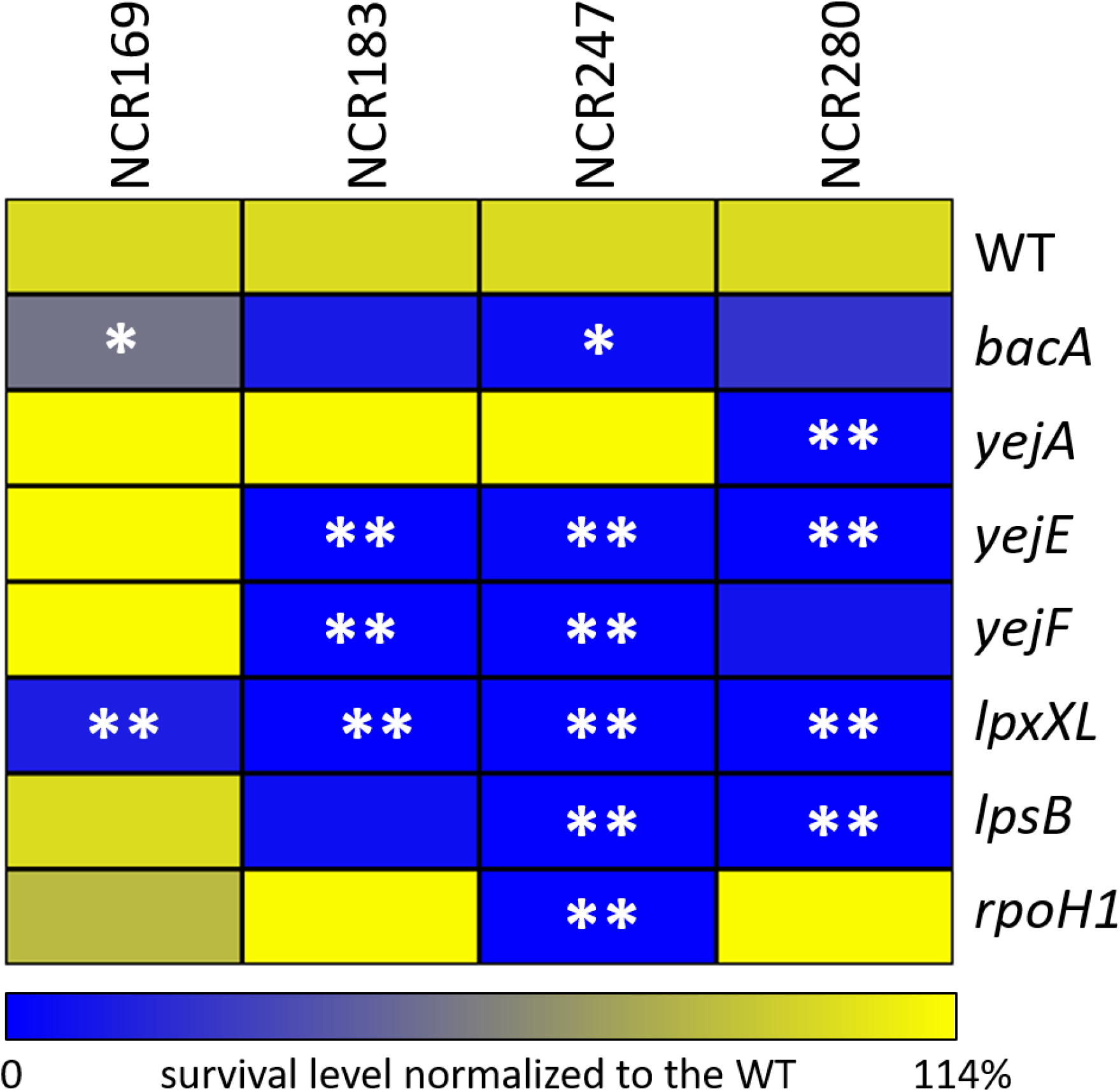
Sensitivity profile of *Sinorhizobium meliloti* strains to a panel of NCR peptides. The heatmap shows the relative survival, expressed in % of the mutant strains compared to the wild type (WT) survival set at 100% for each peptide treatment. Nonparametric Kruskal-Wallis and post-hoc Dunn tests were used to assess the significance of differences which are indicated by asterisks (*\*\** is *p*<0.05; * is *p*<0.1). One representative experiment out of two is shown.

Besides the exposure to the NCR peptides, inside the nodules bacteroids also experience additional stress factors such as elevated hydrogen peroxide levels (34), an acidic pH formed in the peribacteroid space (35) and a microaerobic environment (36). Almost all the mutants behaved very similar to wild type during unstressed growth in culture or in response to hydrogen peroxide, acid, and low oxygen stress (Fig. S2). Only the *rpoH1* mutant was much less resistant to acid and anaerobic stress and had a reduced growth rate (Fig. S2). On the other hand, the mutants were more sensitive than wild type to the exposure to the detergent sodium dodecyl sulfate (SDS) (Fig. S2), indicating that these mutants have a reduced ability to cope with membrane permeabilizing stresses, in agreement with their increased NCR sensitivity. Thus, except for the *rpoH1* mutant, which has a general defect in growth and stress management, the selected mutants seem to be, among the known stress factors in the nodule environment, specifically sensitive to the NCRs.

### Nodule formation by NCR-sensitive *Sinorhizobium meliloti* mutants

Next, the phenotype of these mutants in symbiosis with *M. truncatula* was compared with the wild-type strain and the *bacA* mutant. Measurement of the nitrogen fixation activity and macroscopic inspection of the root system of plants inoculated with the wild type and the seven mutants (Fig. S3) revealed that, besides the wild type, mutants in the *yejA, yejE, yejF* and *lpxXL* genes formed functional nodules (Fix^+^), although the *yejE, yejF* and *lpxXL* mutants had a reduced nitrogen-fixation activity. On the other hand, the *bacA, lpsB* and *rpoH* mutants formed non-functional (Fix^-^) and abnormally looking nodules that were small and white, in agreement with previous descriptions (19, 29, 32).

The histological organization of the nodules formed by the mutants, the formation of infected symbiotic cells and the viability of the bacteria they contain, were analysed using confocal microscopy (Fig. 2A). The used staining procedure highlights the nodule bacteria with a green fluorescence signal (SYTO ™ 9) when their membranes are well preserved and with a red fluorescence (propidium iodide) when their membranes are highly permeable. As previously reported, wild-type nodules formed symbiotic cells infected with green-labelled elongated bacteroids, while the nodules infected with the *bacA* mutant contained symbiotic cells carrying small undifferentiated bacteria stained red (19).

**Figure 2.**
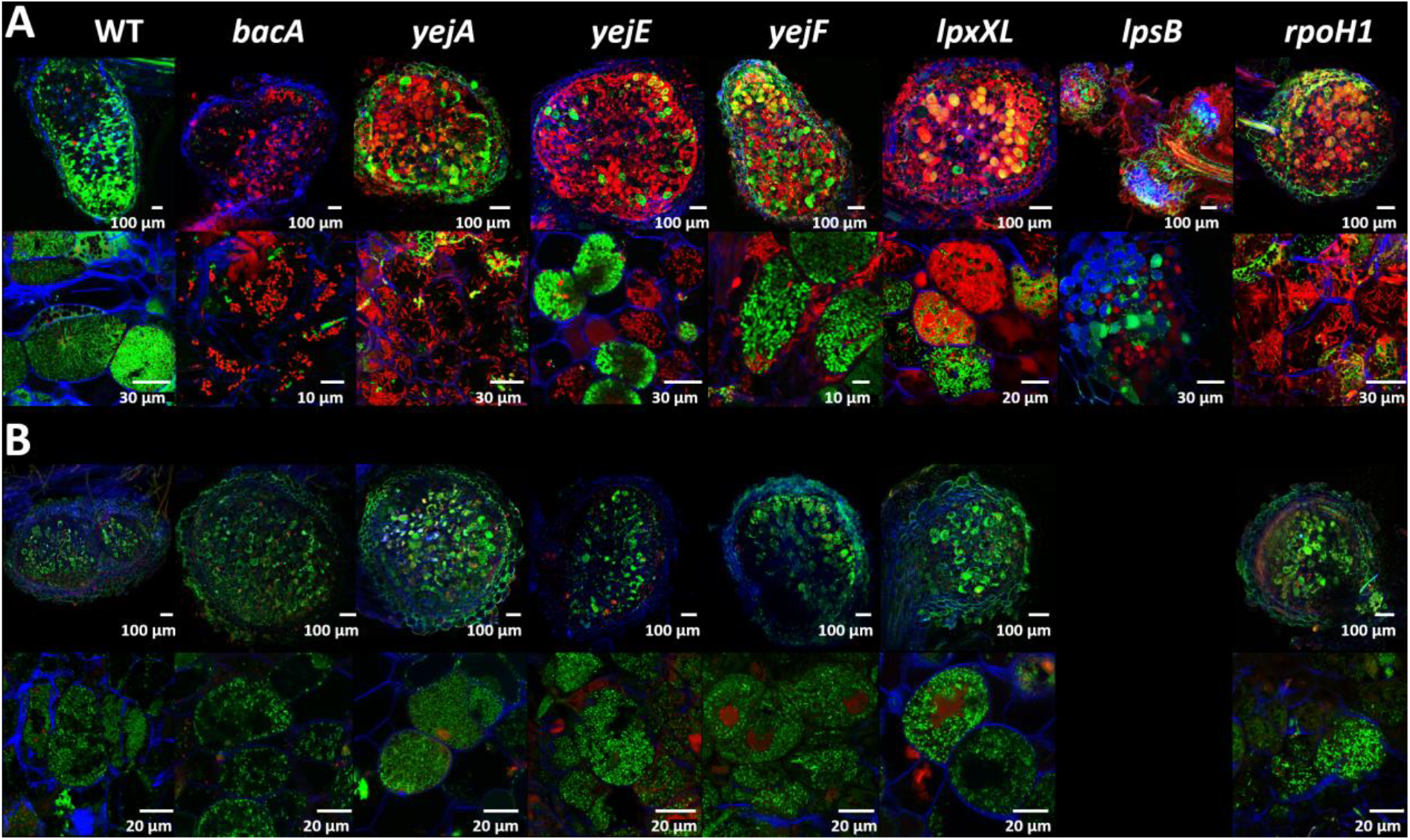
Symbiotic phenotype of *Sinorhizobium meliloti* mutants during symbiosis with *Medicago truncatula* wild type and *dnf1* mutant. **A**. Membrane permeability in bacteroids of *S. meliloti* Sm1021 wild type (WT) and mutants in wild-type *M. truncatula* nodules, determined by live-dead staining of nodule sections and confocal microscopy. **B**. Membrane permeability of *S. meliloti* Sm1021 WT and mutants in nodules of the *M. truncatula dnf1* mutant, determined by live-dead staining of nodule sections and confocal microscopy. Nodule phenotype at 21 days post inoculation. Top row images, full nodule sections; Bottom row images, enlarged images of symbiotic cells. Scale bars are indicated in each panel. The staining procedure of nodule sections with a mixture of the dyes propidium iodide, SYTO ™ 9 and calcofluor-white highlights the bacteroids and nodule bacteria with a green fluorescence signal (SYTO ™ 9) when their membranes are well preserved and with a red fluorescence (propidium iodide) when their membranes are highly permeable. Plant cell walls are stained blue (calcofluor-white). Representative images are shown out of five to 10 nodules analyzed per condition and originating from at least two independent nodulation assays.

Contrary to what we expected from the macroscopic inspection of the nodules and their ability to fix nitrogen, the *yejA, yejE* and *yejF* mutant bacteroids were substantially altered compared to wild-type bacteroids. A high proportion of them were red-stained indicating that their membranes were strongly permeabilized. Nevertheless, other host cells contained bacteroids stained in green by the SYTO ™ 9 dye. LpxXL is known to be important, but not essential to *S. meliloti* during symbiosis with alfalfa (*Medicago sativa*) (37). The mutant forms hypertrophied bacteroids (30). In our experiments, the *lpxXL* mutant displayed elongated bacteroids but mostly permeable to propidium iodide (stained in red). This observation confirmed that *S. meliloti* LpxXL is also essential for the normal bacteroid differentiation process in *M. truncatula* but that it is not crucial for infection and nitrogen fixation. The *rpoH1* mutant formed elongated bacteroids that nevertheless were strongly stained by propidium iodide, in agreement with the Fix^-^ phenotype of the nodules and the described phenotype of the mutant in alfalfa (32). Inside the small bumps elicited by the *lpsB* mutant, no cell seems to be colonised by bacteria as revealed by confocal microscopy. Thus, the *S. meliloti lpsB* mutant failed to colonize the host cells and was defective at a stage before plant cell infection. It was previously reported that the *lpsB* mutant colonizes nodules cells in *M. sativa* suggesting a more severe defect in *M. truncatula* (38, 39). Therefore, we tested the phenotype of the selected mutants also on *M. sativa* (Fig. S4). The *lpsB, rpoH1* and the *yejA* mutants had indeed milder defects on this host while the other mutants had similar phenotypes on the two hosts.

### Can the increased membrane permeability of the NCR-sensitive *Sinorhizobium meliloti* mutants be attributed to the NCRs in nodules?

The high membrane permeability of the *bacA* mutant nodule bacteria is the result of the action of the NCR peptides in the nodule cells on this NCR-hypersensitive mutant (19). Possibly, the same is true for the other mutants. To test this hypothesis, we made use of the *M. truncatula dnf1* mutant, which is defective in a nodule-specific subunit of the signal peptidase complex (40). This mutant cannot transport NCR peptides to the symbiosomes (13). As a result, the wild-type *S. meliloti* nodule bacteria in this mutant are not differentiated. The *bacA* mutant, which is strongly permeabilized and stained by propidium iodide in nodules of wild-type *M. truncatula* plants, is not so in the *dnf1* mutant nodule cells because the bacteria are not challenged anymore with the NCRs (Fig. 2B)(19). Similarly to the *bacA* mutant, the *yejA, yejE, yejF, lpxXL* and *rpoH1* mutants did not display membrane permeability, as revealed by the absence of propidium iodide staining, in the infected nodule cells of the *dnf1* mutant (Fig. 2B). This thus suggests that these mutants become membrane-permeabilized by the action of the NCRs and that their symbiotic defects are at least in part due to their hypersensitivity to the NCRs. The *lpsB* mutant did not form detectable nodule-like structures on the *dnf1* roots. Therefore, we cannot conclude on the involvement of the NCR peptides in the symbiotic phenotype of this mutant.

### Bacteroid differentiation of the NCR-sensitive *Sinorhizobium meliloti* mutants

A very strong cell enlargement and an increase in the ploidy level of the bacteria characterize the differentiated bacteroids in *Medicago* nodules. These parameters can be readily measured by DAPI staining and flow cytometry (10). The wild-type bacteroids from *M. truncatula* nodules had a high DNA content, over twenty fold higher than the DNA content in free-living *S. meliloti* (Fig. 3) and increased light-scattering parameters reflecting the cell enlargement (Fig. S5). As previously reported, the nodules infected by the *bacA* mutant did not contain differentiated bacteria (Fig. 3, Fig. S5)(21). The bacteria in nodules induced by the *lpsB* mutant, probably located in infection threads, had a profile that confirmed the complete absence of differentiated bacteria (Fig. 3, Fig. S5). Nodules infected with the *rpoH1* mutant had mostly undifferentiated bacteria although a small amount of fully differentiated cells was detected, as well as cells in an intermediate stage. Also the nodules of the *lpxXL* mutant contained many undifferentiated bacteria as well as fully differentiated ones (Fig. 3, Fig. S5). In contrast, the *yejA, yejE* and *yejF* mutant nodules contained large numbers of fully differentiated bacteria (Fig. 3, Fig. S5).

**Figure 3.**
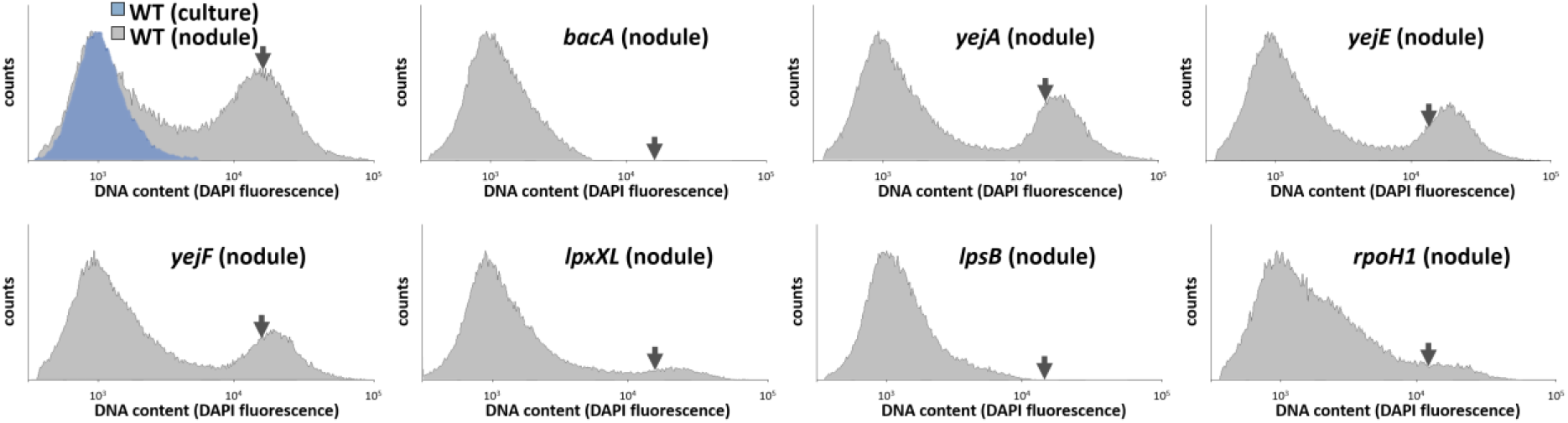
DNA content of nodule bacteria in *Medicago truncatula*. Flow cytometry analysis of the DNA content of bacteria in culture or isolated from nodules infected with the indicated strains and stained with 4’,6-diamidino-2-phenylindole (DAPI). The cell counts (y-axes) are represented in function of the DAPI fluorescence intensity (x-axes). The arrow in each graph indicates the mean DNA content of wild-type (WT) bacteroids as in the upper left panel. One representative experiment out of two is shown.

It was reported previously that the *lpxXL* mutant formed nodules containing hypertrophied and larger bacteria than the wild-type bacteroids (30). In our study, this difference was detectable in the flow cytometry measurements showing a higher DAPI fluorescence and light scattering for the small portion of differentiated bacteria in these nodules (Fig. 3, Fig. S5). Moreover, we noticed that the bacteroids in the nodules infected with the *yejA, yejE* and *yejF* mutants displayed similar higher DNA fluorescence and light scattering (Fig. 3, Fig. S5), suggesting that also these bacteroids have abnormal morphologies.

These patterns in *M. truncatula* nodules were overall similar in *M. sativa* although the *rpoH1* mutant showed a higher number of intermediate and fully differentiated bacteroids and the *yejA* mutant had a profile similar to the wild type in nodules at 21 dpi while larger than normal bacteroids were only detected at 32 dpi (Fig. S6). These differences corresponded well with the macroscopic and microscopic differences in the nodules between the two host plants (Fig. 2, Fig. S3, Fig. S4).

### Defective bacteroid differentiation of *yejE* and *yejF* mutants

To confirm the altered bacteroid morphologies of the *yej* mutants suggested by the cytometry analysis, the *yejE* and *yejF* mutants were observed at high magnification by microscopy. Confocal and transmission electron microscopy of *M. sativa* nodule sections showed that the nodule cells infected with the *yejE* or *yejF* mutant contained a very heterogeneous population of abnormal bacteroid morphs, including elongated, spherical, club-shaped or irregular blob-like cells (Fig. 4A, Fig. S7). These cells contrasted strongly with the narrow, elongated wild-type bacteroids. This aberrant bacteroid phenotype of the mutants was restored to a wild-type phenotype by the introduction of a plasmid-borne copy of the *yejABEF* genes (Fig. S7). The difference between wild-type and mutant bacteroids was also obvious by fluorescence microscopy observations of purified nodule bacteria (Fig. 4B). The quantification of cell morphology parameters by image analysis with MicrobeJ of free-living bacteria and the purified nodule bacteria confirmed the irregular forms of *yejE* and *yejF* bacteroids, notably showing that these bacteroids are broader and more spherical than the elongated wild-type bacteroids (Fig. 4C). Transmission electron microscopy further showed that the cytoplasm and inner membrane of many of the *yejF* bacteroids was retracted, leaving very large intermembrane spaces, which in some cases even developed into vacuoles, entirely surrounded by cytoplasm. Some cells had multiple small and large vacuoles (Fig. S7A). In contrast to the bacteroids, in the cultured bacteria, no differences were observed in the ultrastructure of the wild-type and *yejF* mutant bacteria (Fig. S7B). The formation of strongly abnormal bacteroids of the *yej* mutants is *a priori* contradictory with the nitrogen fixation activity of these nodules. Expression of the *nifH* gene is a marker for nitrogen-fixing bacteroids (41). A *GFP* gene under the control of the *nifH* promoter was introduced into the wild-type strain and the *bacA* (as a negative control), *yejE* and *yejF* mutants. Analysis of the GFP fluorescence in nodule bacteria of *M. truncatula* (Fig. 5) and *M. sativa* (Fig. S8) by confocal microscopy and flow cytometry showed that despite their aberrant morphology, the *yejE* and *yejF* bacteroids are mostly functional, although some nodule cells contain highly permeable bacteroids, lacking the GFP-signal suggesting that they were non-functional.

**Figure 4.**
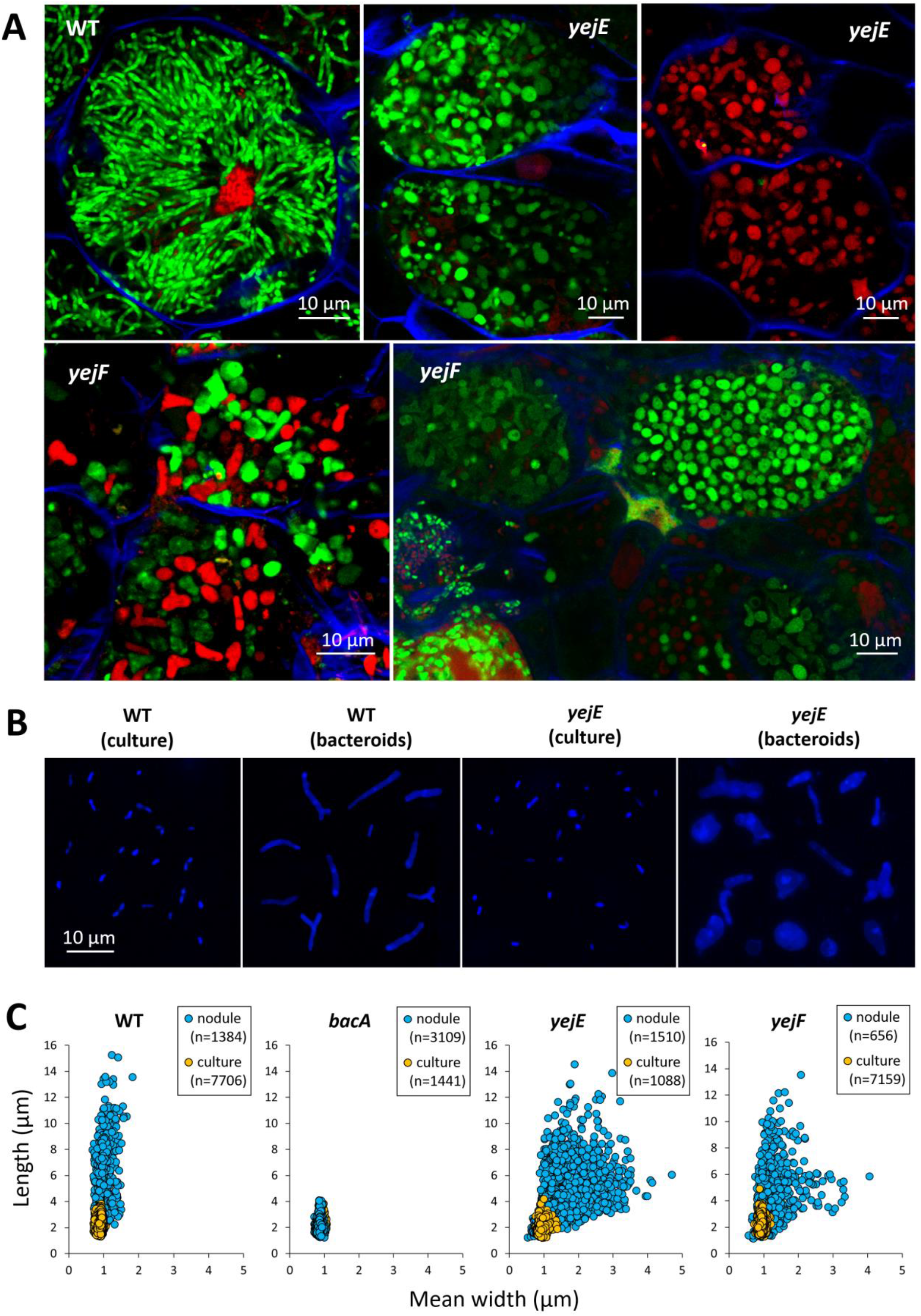
Bacteroid morphology of *yejE* and *yejF* mutants in *Medicago sativa* nodules. **A**. Sections of nodules infected with the wild type (WT) and the *yejE* or *yejF* mutants were stained with a mixture of the dyes propidium iodide, SYTO ™ 9 and calcofluor-white and observed by confocal microscopy. Scale bars (10 µm) are indicated in each panel. Representative images are shown out of five to 10 nodules analyzed per condition and originating from at least two independent nodulation assays. **B**. Preparations of cultured bacteria or purified *M. sativa* nodule bacteria of the wild type and the *yejF* mutant were observed by fluorescence microscopy. The panels are composite images and the shown individual cells were cut from original images and recombined in a single panel. Each panel is at the same magnification and the scale bar (10 µm) is indicated in the left panel. **C**. Cell shape parameters of free-living bacteria and bacteroids determined by MicrobeJ. Dot plots show the mean width and length of cells of the indicated strains. Each point represents the values of one bacterium. Numbers of analysed cells (n) are indicated.

**Figure 5.**
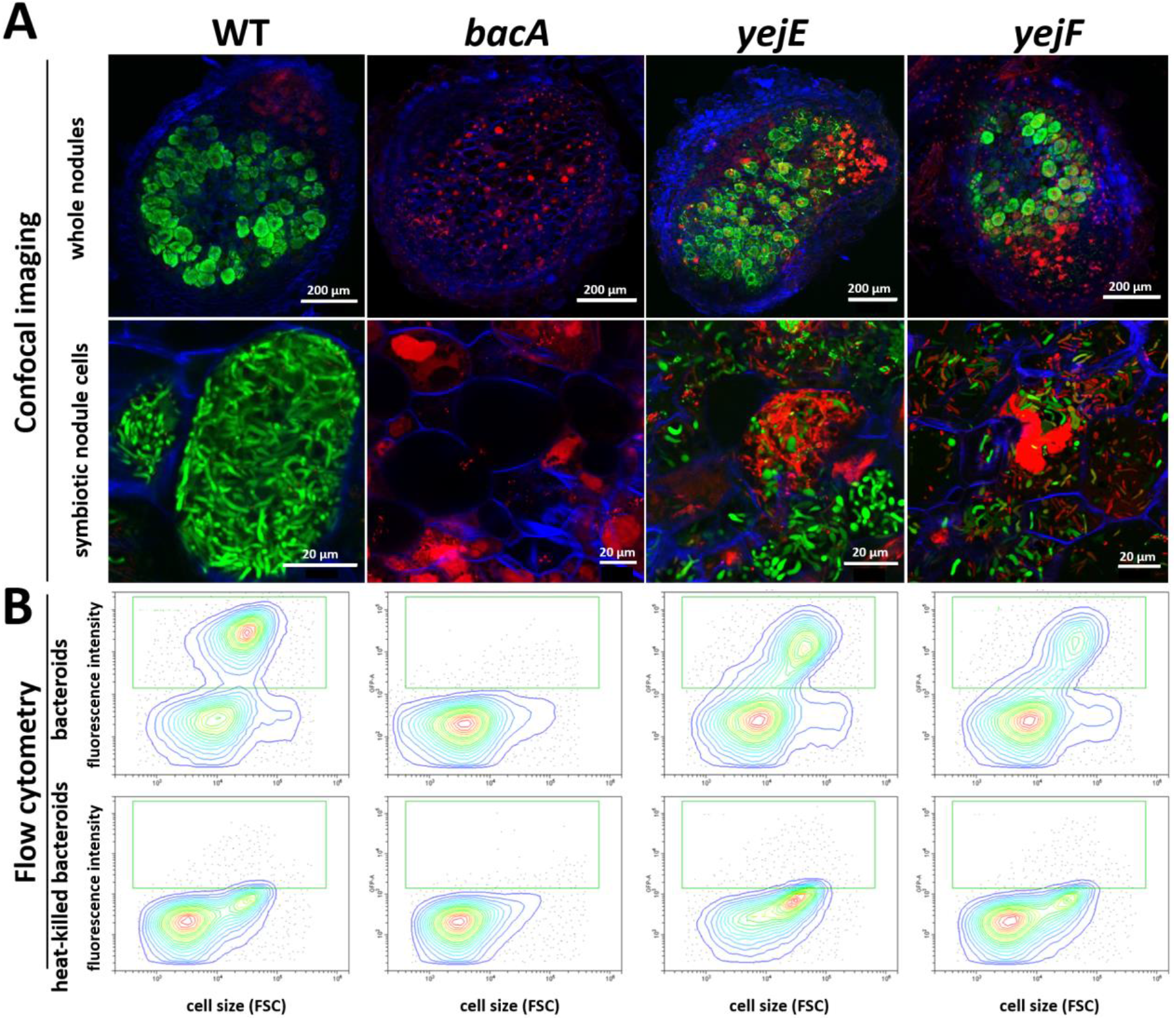
Nitrogenase expression in the *yejE* and *yejF* mutant bacteroids in *Medicago truncatula* nodules. **A**. Confocal microscopy of sections of nodules infected with *S. meliloti* Sm1021.p*pnifH*-*GFP* (WT), *bacA*.p*pnifH*-*GFP* (*bacA*), *yejE*.p*pnifH*-*GFP* (*yejE*) or *yejF*.p*pnifH*-*GFP* (*yejF*) and stained with propidium iodide (red stain). Green stained bacteroids are functional while red stained bacteroids are non-functional. **B**. Flow cytometry determination of GFP levels in nodule bacteria (upper panels) and heat-killed nodule bacteria (lower panels). The green square shows the position of the GFP-positive bacteroids. FSC is forward scatter.

### NCR247 uptake by the YejABEF transporter

In *Escherichia coli*, both YejABEF and the BacA homolog SbmA mediate the transport of microcin C peptide-nucleotide antibiotics. Therefore, we tested whether the overlap in substrates of YejABEF and BacA can be extended to NCR peptides, which are known substrates of BacA (20, 21). We tested the impact of the mutations in the YejABEF transporter genes on NCR247 uptake and compared it with a mutation in *bacA*. With a flow cytometry based assay and a fluorescent derivative of the NCR247 peptide, we found that while the NCR247 uptake is completely abolished in the *bacA* mutant, as expected, its uptake is also reduced but not completely abolished in the *yejE* and *yejF* mutants (Fig. 6). This suggests that the YejABEF transporter contributes to NCR uptake.

**Figure 6.**
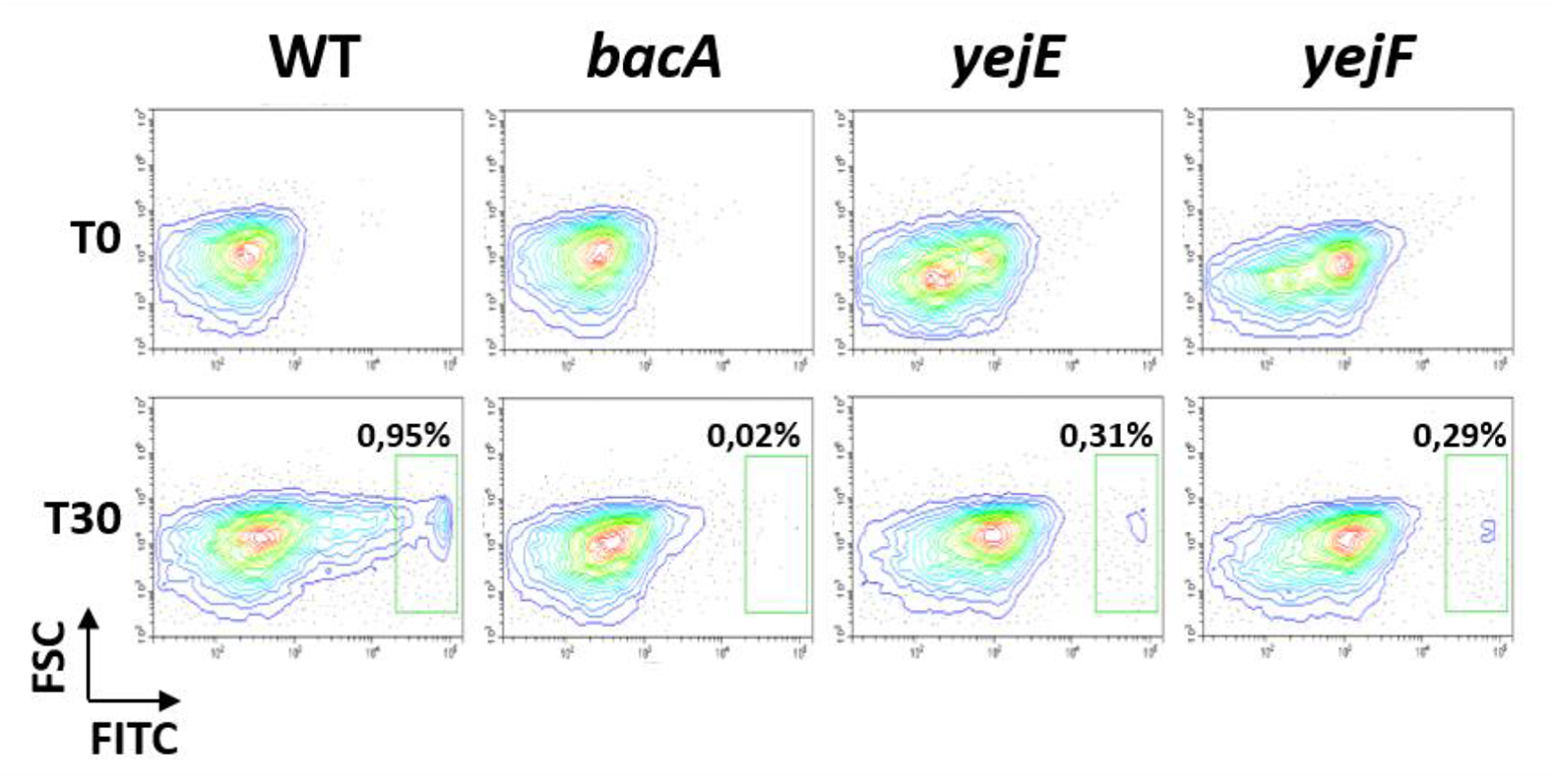
Uptake of NCR247 mediated by the BacA and YejABEF transporters. Flow cytometry measurement of FITC fluorescence in *S. meliloti* cells before (T0) and after 30 minutes of incubation (T30) of the bacteria with the NCR247-FITC peptide. FITC in the x-axis is FITC-derived fluorescence and FSC in the y-axis is forward scatter. The green square shows the bacteria that are FITC-positive and have uptaken the NCR247-FITC peptide. In the wild type (WT), a subpopulation of the bacteria import the NCR247-FITC peptide and in the *bacA* mutant, this population is absent. In the *yejE* and *yejF* mutants, the FITC-positive subpopulation is strongly reduced compared to wild type. The analysis was performed in quadruplicate and a representative example is shown.

## Discussion

### Multiple functions of *Sinorhizobium meliloti* contribute to NCR resistance and are required for bacteroid formation and persistence

The NCR peptides are a two-edged sword. On the one hand, they maneuver the rhizobial endosymbionts into a terminally differentiated state via a multitude of activities on the bacteria from which the membrane permeabilization, cell cycle perturbation (polyploidization) and cell enlargement are the most visible ones. On the other hand, many NCRs have antimicrobial activity, notably the cationic ones, potentially killing the endosymbionts (13, 14). Therefore, rhizobia have to defend themselves to be able to establish a chronic infection in the NCR-producing symbiotic cells of the nodules. Previous work has identified the BacA peptide transporter and the EPS barrier in the bacterial envelope as defenses of *Sinorhizobium* strains against the NCRs of their *Medicago* hosts (14, 19, 23).

Here, we defined three new functions in *S. meliloti*, the LPS, the sigma factor RpoH1, and the YejABEF peptide transporter as additional determinants in bacteroids, required to cope with the NCR peptides. We show that knockout mutants in the corresponding genes are more sensitive to a panel of antimicrobial NCRs and that this hypersensitivity is correlated with a strongly enhanced membrane permeability of the nodule bacteria and abnormalities in their morphology and ploidy levels. It is striking, however, that the different mutants have markedly different bacteroid phenotypes, ranging from undifferentiated to hyper-differentiated. This can be attributed to at least three factors. Each mutant has a specific NCR-sensitivity profile when tested against a small panel of peptides and the “NCR landscape” present in the developing symbiotic cells is continuously changing because of the expression of the *NCR* genes in different waves during symbiotic cell differentiation (15). Accordingly, each mutant could accumulate NCR-induced damage at a different rate and reach the breaking point at different stages over the course of the bacteroid differentiation process (Fig. S1). Moreover, the expression pattern of the bacterial genes in the nodule zones is different, suggesting that their principal impact is realized at distinct stages of the symbiotic cell development and bacteroid differentiation.

Blocking the targeting of the NCR peptides to the nodule bacteria by the use of the *M. truncatula dnf1* mutant prevents the membrane permeabilization of the nodule bacteria in the mutants. This observation places the symbiotic role of these bacterial genes downstream of the peptide targeting to the symbiosomes (except for *lpsB*, see below) and links them, at least in part, to the NCRs. It should be noted that besides the antimicrobial NCR peptides, the other stress factors present in the nodule environment, such as high H_2_O_2_ levels, a low pH and low oxygen (34-36), are possibly also reduced or absent in the *dnf1* nodules since they are non-functional (40). However, we found that, except for *rpoH1*, mutants in the studied genes were not differently affected than the wild type by these stresses, making it unlikely that they are the main cause of the bacteroid phenotypes in these mutants.

The functions of *S. meliloti* described here and before probably picture only part of the full toolkit of this symbiont to survive the NCR challenge. Many additional *S. meliloti* mutants are described with symbiotic phenotypes that are suggestive for a similar contribution to NCR resistance. (26, 42–53). In addition, other genes that could be important for symbiosis, were discovered in a Tn-seq screen with the peptide NCR247 (25). This screen identified here-described functions like the BacA and YejABEF transporters, the LPS and EPS biosynthesis, but also never analyzed functions. It would be of major interest to explore the function of these genes in relation to the NCR response of bacteroids. Note that some of these genes might have escaped identification in previous genetic screens for symbiosis mutants because mutations might provoke only subtle phenotypes with a weak overall effect on nitrogen fixation in standard laboratory conditions but with bacteroid alterations in morphology and persistence as illustrated here with the *yejABEF* mutants.

### Response to NCR-induced stress regulated by the alternative sigma factor RpoH1

Bacteria deal with different types of stress conditions by global transcriptional responses mediated by alternative sigma factors. As a general classification, the RpoH sigma factors are believed to control cytoplasmic stress responses, while the RpoE sigma factors are known to respond to periplasmic and membrane stress (54). Given the membrane damage provoked by NCRs and the general function of RpoE sigma factors in envelope stress response, including in response to AMPs (55), a role of the RpoE regulators of *S. meliloti* in bacteroid differentiation and NCR response could *a priori* have been expected. *S. meliloti* has 11 RpoE-like sigma factors. Remarkably, despite the fact that some RpoEs have a considerable effect on gene transcription, all single mutants, all possible double mutants and even a mutant lacking all 11 genes showed no detectable phenotypic difference with wild type in symbiosis or during many tested free-living growth conditions, including growth in the presence of membrane stresses (56, 57). Thus, the 11 *S. meliloti* RpoEs do not have the expected role in regulating the envelope stress response.

This role seems to be taken up in part by RpoH in *S. meliloti*, in agreement with its role in symbiosis and NCR resistance and the observation that growth of the *rpoH1* mutant is affected in the presence of various membrane-disrupting agents (32). The latter suggests that RpoH in *S. meliloti* is not respecting the canonical division of labor between alternative sigma factors and that its function is not restricted to cytoplasmic stress, including acidic stress and anoxia or microoxia, but also encompasses membrane stresses. We propose that the here uncovered role of RpoH1 in NCR resistance and in bacteroid differentiation is connected to its regulation of membrane stress.

### The lipopolysaccharide barrier against NCR membrane damage

In Gram-negative bacteria, including rhizobia, LPS constitutes the first point of attack of cationic AMPs. In a two-stage process, AMPs make first electrostatic interactions with the negatively charged LPS, allowing the AMPs to approach the membrane lipids and subsequently to insert into the lipid bilayer, perforate it and translocate into the periplasm (58). The chemical composition of LPS can influence the efficiency of the AMP attack and pathogens have evolved mechanisms to recognize the presence of AMPs and to modify in response the composition of the LPS to lower the potency of the AMP attack (59).

*S. meliloti lpsB* encodes a glycosyl transferase that participates in the biosynthesis of the LPS core (60). The mutation of the gene affects both the core structure and O-antigen polymerization (41, 42). The O-antigen is thought to be a camouflage, masking the membrane and the charges in the membrane vicinity. In the *lpsB* mutant, the shorter O-antigen could be a less efficient shield against the NCRs, offering to the peptides an easier access to membrane-proximal charges resulting in the increased sensitivity of the *lpsB* mutant to the NCRs (this work) as well as to other AMPs (29, 39). The *lpsB* mutant forms nodules on *M. sativa*, which contain infected cells and it has a phenotype that is similar to the *bacA* mutant. However, the *lpsB* mutant forms uninfected nodules on *M. truncatula* roots. Therefore, it was not possible to test the implication of NCRs in this symbiotic phenotype with the use of the *dnf1* mutant, since the *DNF1* gene is expressed only in infected nodule cells (40). Since the *NCR* genes are nearly exclusively expressed in the infected symbiotic cells (15), the blockage of the mutant before the release of bacteria into nodule cells suggests that the LPS could provide protection against another stressor produced very early on in the infection process in *M. truncatula*. On the other hand, it was recently reported that some *NCR* genes are expressed in infected root hairs or Nod factor-stimulated root epidermal cells (61, 62). Thus, it is possible that the challenge with these early NCR peptides is already detrimental to the mutant, blocking any further progress in the infection process.

LpxXL is a specific acyl transferase that introduces in the lipid A moiety of LPS the very long chain fatty acid 27-OHC28:0 (37). This LpxXL-dependent acylation is expected to make the lipid A hydrophobic, forming the biophysical basis for LpxXL-dependent NCR resistance. Hydrophobic lipid A increases the thickness of the outer layer of the outer membrane and reduces the membrane fluidity, which on its turn prevents or delays AMP insertion and membrane damage. Increasing lipid A hydrophobicity by introducing additional acyl chains is a well-known mechanism used by *Salmonella* to enhance its resistance against host AMPs during infection (59).

### The YejABEF peptide transporter provides resistance to membrane-damaging peptides and sensitizes bacteria to AMPs with intracellular targets

The YejABEF ABC transporter has never been analyzed before in the context of the rhizobium-legume symbiosis. Furthermore, even though this transporter is highly conserved among proteobacteria, its physiological role in whichever bacterium has been characterized only in a few instances. One of them is the uptake of microcin C in *E. coli*. This translation-inhibiting peptide-nucleotide of bacterial origin has no action on the bacterial membrane but has an intracellular target, the Asp aminoacyl-tRNA synthase (63). A mutant in the YejABEF transporter cannot uptake microcin C and is resistant to it (64, 65). Conversely, the *Salmonella* and *Brucella yejABEF* mutants are more sensitive to peptides with membrane-damaging activity such as defensins, polymyxin B, protamine and melittin. Consequently, these mutants have reduced pathogenicity because their capacity to survive in macrophage cell lines or in mice is diminished (66, 67). Thus, the increased sensitivity of the *yejA, yejE* and *yejF* mutants of *S. meliloti* towards NCRs is consistent with these previous findings.

The characteristics of the YejABEF peptide transporter towards membrane-damaging peptides versus peptides with intracellular action are intriguingly parallel to the features of the BacA peptide transporter (SbmA in *E. coli* and *Salmonella*). These latter transporters are required for the import of diverse peptides with intracellular targets (microcins B17 and J25, Bac7, Bac5 and bleomycin) and mutants are therefore resistant to them while the same mutants are hypersensitive to membrane-active peptides (defensins, NCRs) (19, 68, 69, 70, 71). Moreover, the range of peptides that can be imported overlap between these two transporters. Both, YejABEF and SbmA can uptake microcin C derivatives (64). Furthermore, we show that both BacA and YejABEF contribute to NCR247 uptake. Intriguingly, our genetic analysis suggests that both transporters cooperate to import this peptide, since inactivation of one or the other abolishes or strongly reduces uptake. How these two transporters can physically interact and cooperate is an issue of interest for future research. Nevertheless, BacA can function (partially) in the absence of YejABEF while the inverse is not the case. This could be the basis for the markedly different symbiotic phenotype of the *bacA* and *yej* mutants. While the *bacA* mutation is detrimental from the earliest contact with NCRs and bacteria die as soon as they are released from infection threads in the symbiotic cells, the *yej* mutants cope with the NCR peptides much longer and show abnormalities only at the end of the bacteroid differentiation process and symbiont lifespan. The *yejA* gene encodes the periplasmic binding protein of the transporter. The *yejA* mutant had a different NCR sensitivity profile than the *yejE* and *yejF* mutant and this was correlated with a different symbiotic phenotype in the *M. sativa* host. This suggest that YejA contributes to the interaction with only a subset of the NCR peptides.

How do the YejABEF and BacA transporters contribute to resistance to NCRs and other membrane-damaging peptides? The most straightforward model that has been proposed before for BacA (19– 21) as well as for the unrelated SapABCDF peptide uptake transporter (72), is the reduction of the AMP concentration in the vicinity of the inner membrane below a critical threshold. Alternatively, the presence or activity of the transporters might indirectly affect the bacterial envelope structure, rendering it more robust against AMPs. The higher sensitivity of the *yej* mutants towards SDS is in agreement with this possibility. Similarly, membrane alterations were reported in the *bacA* mutant of *S. meliloti* (73).

## Conclusions

The multifaceted NCR resistance required for symbiosis and chronic infection of the nodule cells mirrors the multitude of AMP resistance mechanisms in animal pathogens, which collectively contribute to the pathogenicity of these bacteria (2, 3, 74). However, one dimension of this strategy in pathogens is not known in *S. meliloti* and consists of the direct recognition of host AMPs by receptors triggering an adaptive response. Probably the best-studied AMP receptor is the two-component regulator PhoPQ in *Salmonella*, which adjusts the LPS composition in response to the presence of host peptides (59). Perhaps the *S. meliloti* ExoS-ChvI or FeuP-FeuQ two-component regulators, which are upregulated by NCR treatment and essential for symbiosis (26, 42, 43), form such a regulatory module, recognizing NCRs and controlling an appropriate response in the symbiotic nodule cells. In this respect, it is of interest to note that the rhizobial LPS structure changes strongly in bacteroids of NCR-producing nodules (75, 76).

As shown here, rhizobia have to defend themselves to be able to establish a chronic infection in the NCR-producing symbiotic cells of the nodules. On the other hand, the profile of NCR peptides produced in the nodule cells is also determinant for the outcome of the symbiosis and some *M. truncatula* mutants in individual *NCR* genes or *M. truncatula* accessions, expressing specific *NCR* alleles, display incompatibility with *S. meliloti* strains (77–80). Thus, a fine balance must be established in the symbiotic nodule cells between the NCR landscapes and matching multifactorial bacterial countermeasures. Perturbations in the host or in the endosymbiont, as the ones described here, affecting this equilibrium lead to a breakdown of the symbiosis.

## Materials and Methods

### Bacterial strains, plant growth and nodulation assays and analysis

The procedures for the growth of the *S. meliloti* Sm1021 strain and its derivatives, plant culture of the *M. sativa* cultivar Gabès, the *M. truncatula* accession Jemalong A17, and the A17 *dnf1* mutant, nodulation assays, acetylene reduction assays, bacteroid isolation, flow cytometry measurements and confocal microscopy were performed as described before (81). Bacterial mutants in *rpoH1, yejA, yejE* and *yejF* were obtained by plasmid insertion and gene deletion (Table S2) as described before (82). A fragment containing the *yejABEF* genes was PCR-amplified from *S. meliloti* Sm1021 genomic DNA using primers yejA_NdeI_F and yejF_XbaI_R (Table S2) and cloned between the NdeI and XbaI restriction sites of the pSRK(Gm) vector (83). The construct was confirmed by restriction analysis and sequencing. The plasmid as well as the empty pSRK(Gm) vector were conjugated to wild type Sm1021 as well as the *yejA, yejE*, and *yejF* mutants bi tri-parental mating (82).

Quantitative analysis of the shape of free-living bacteria and bacteroids was performed with the MicrobeJ plugin of ImageJ (84). Bacteroid extracts and exponential phase cultures were stained with 2.5 nM SYTO ™ 9 for 10 minutes at 37°C and mounted between slide and coverslip. Bacteria imaging was performed on a SP8 laser scanning confocal microscope (Leica microsystems) equipped with hybrid detectors and a 63x oil immersion objective (Plan Apo, NA: 1.4, Leica). For each condition, multiple z-stacks (2.7 µm width, 0.7 µm step) were automatically acquired (excitation: 488 nm; collection of fluorescence: 520-580 nm). Stacks were transformed as maximum intensity projections using ImageJ software (85). Bacteria in the image stacks were automatically detected with MicrobeJ using an intensity based threshold method with a combination of morphological filters. To ensure high data quality every image was manually checked to remove false positive objects (mainly plant residues). Morphological parameters were directly extracted from MicrobeJ and figures were created with Excel or ggplot2 in R.

### *In vitro* sensitivity and peptide uptake assays

NCR sensitivity assays were carried out essentially as described (20). Triplicate measurements were performed in at least two independent experiments per treatment. Nonparametric Kruskal-Wallis and post-hoc Dunn tests were performed to assess significance of differences between the sensitivity of the wild type and the mutant strains. To measure the resistance of strains to sodium dodecyl sulfate (SDS), H_2_O_2_ and HCl stress, overnight cultures of the wild type and mutants were diluted to OD_600_=0.2. A total of 100 μL of these suspensions was added to 3 ml soft agar (0.7% agar) and poured onto standard 1.5% agar plates. After solidification of the soft agar, filter paper disks (5 mm diameter) were placed on the center of the plate, and 5 μL of 10% (w/v) SDS, 2 M HCl or 1% H_2_O_2_ was added to the disks. Plates were incubated at 28°C for 3 days, and the diameter of the clearing zone was measured. For microaerobic and anaerobic treatments, cultures of the wild type and mutants were diluted to OD_600_=0.1 and 5-fold dilution series were prepared of each. The dilution series were subsequently spotted (5 µL per spot) on agar plates. One series of plates was grown in standard aerobic conditions. A second series of plates was placed in a 2.5 L airtight jar containing an AnaeroPack-MicroAero (Mitsubishi Gas Chemical) bag for microaerobic growth (6-12% O_2_ according to the product specifications). The third series of plates was placed in a 2,5 L jar containing an AnaeroGen 2,5L (Thermo Scientific) bag for anaerobic growth (<0,1% O_2_ according to the product specifications). All plates were grown at 28°C and after 3 days, the plates were removed from the jars for the microaerobic and anaerobic conditions. After 3 days, colonies were sufficiently grown in the aerobic condition for counting. Growth was observed in the microaerobic condition but less strong than in the aerobic condition and these plates were further incubated for 1 day before colony counting. No growth was observed after 3 days in the anaerobic condition and plates were incubated for 4 days in aerobic condition before colony counting. Colony counting was done using a binocular. Quadruplate measurements and two independent experiments per treatment were performed. Nonparametric Kruskal-Wallis and post-hoc Dunn tests were performed to assess significance of differences in all the assays. For growth curves, precultures were diluted to OD_600_=0.04 in YEB medium (81). Cultures were dispatched in microtiter plates with 200 µL of culture per well. The plates were incubated in a SPECTROstar Nano (BMG Labtech) plate reader during 100 h, at 28°C, 100 rpm shaking and OD_600_ measurements every 30 min. Doubling times were calculated from the obtained growth curves (n=10) and nonparametric Kruskal-Wallis and post-hoc Dunn tests were performed to assess significance of differences between growth rates. Peptide uptake assays were performed as described in quadruplicate (20, 86).

### TEM

Bacterial suspensions at OD_600_=6 or 21 days old nodule samples were incubated in fixative (3% glutaraldehyde, 1% paraformaldehyde in 0.1 M cacodylate, pH 6.8) for one to three hours, and washed with 0.1 M cacodylate buffer, pH 6.8. Samples were then incubated one hour in 1% osmium tetroxide, 1.5% potassium ferrocyanide in water. After washing, bacterial suspensions were pelleted in 2% low melting point agarose to facilitate their manipulation. Samples were dehydrated by incubation in increasing concentrations of ethanol (nodule samples for a total of 4 hours in 10-20-30-50-70-90-100%-absolute ethanol-propylene oxide; bacterial pellets for a total of 2 hours in 10-30-50-70-90-100%-absolute ethanol), followed by infiltration with epoxy resin (low viscosity Premix Kit medium, Agar) (for nodule samples, 3 days of infiltration; for bacterial pellets, 24h of infiltration) and polymerization for 24h at 60°C. Ultrathin sections (80-70 nm) were obtained with an ultramicrotome EM UC6 (Leica Microsystems) and collected on formvar carbon-coated copper grids (Agar). Sections were stained with 2% uranyl acetate (Merck) and lead citrate (Agar) before observation with a JEOL JEM-1400 transmission electron microscope operating at 120kV. Images were acquired using a postcolumn high-resolution (11 megapixels) high-speed camera (SC1000 Orius; Gatan) and processed with Digital Micrograph (Gatan).

## Acknowledgements

We are grateful to Corinne Foucault for assistance with plant experiments and Cynthia Dupas for help with the TEM experiments. QN and SD were supported by PhD fellowships from the Paris-Saclay University and QB and NB by Post-doc grants from the Agence Nationale de la Recherche. The present work has benefited from Imagerie-Gif core facility supported by the Agence Nationale de la Recherche (ANR-11-EQPX-0029/Morphoscope, ANR-10-INBS-04/FranceBioImaging; ANR-11-IDEX-0003-02/Saclay Plant Sciences). The EK laboratory is supported by the NKFIH Frontline Research project KKP129924 and a Balzan research grant. The work in the PM laboratory was supported by the LabEx Saclay Plant Sciences-SPS and grants ANR-17-CE20-0011 and ANR-16-CE20-0013 from the Agence Nationale de la Recherche.

## Author contributions

QN, QB and NB have initiated jointly this work. QN, QB, NB, SD, DT, ML, EB and TT constructed strains and performed nodulation assays. QN, QB, SD, RLB, CB and TT performed microscopy and flow cytometry. QB, SJ, AK and EK performed the *in vitro* experiments and provided reagents. QN, QB, NB, EB, TT, BA and PM conceived the study and analyzed the data. PM wrote the manuscript with input from all authors.

## Figures and legends

**Figure S1.**
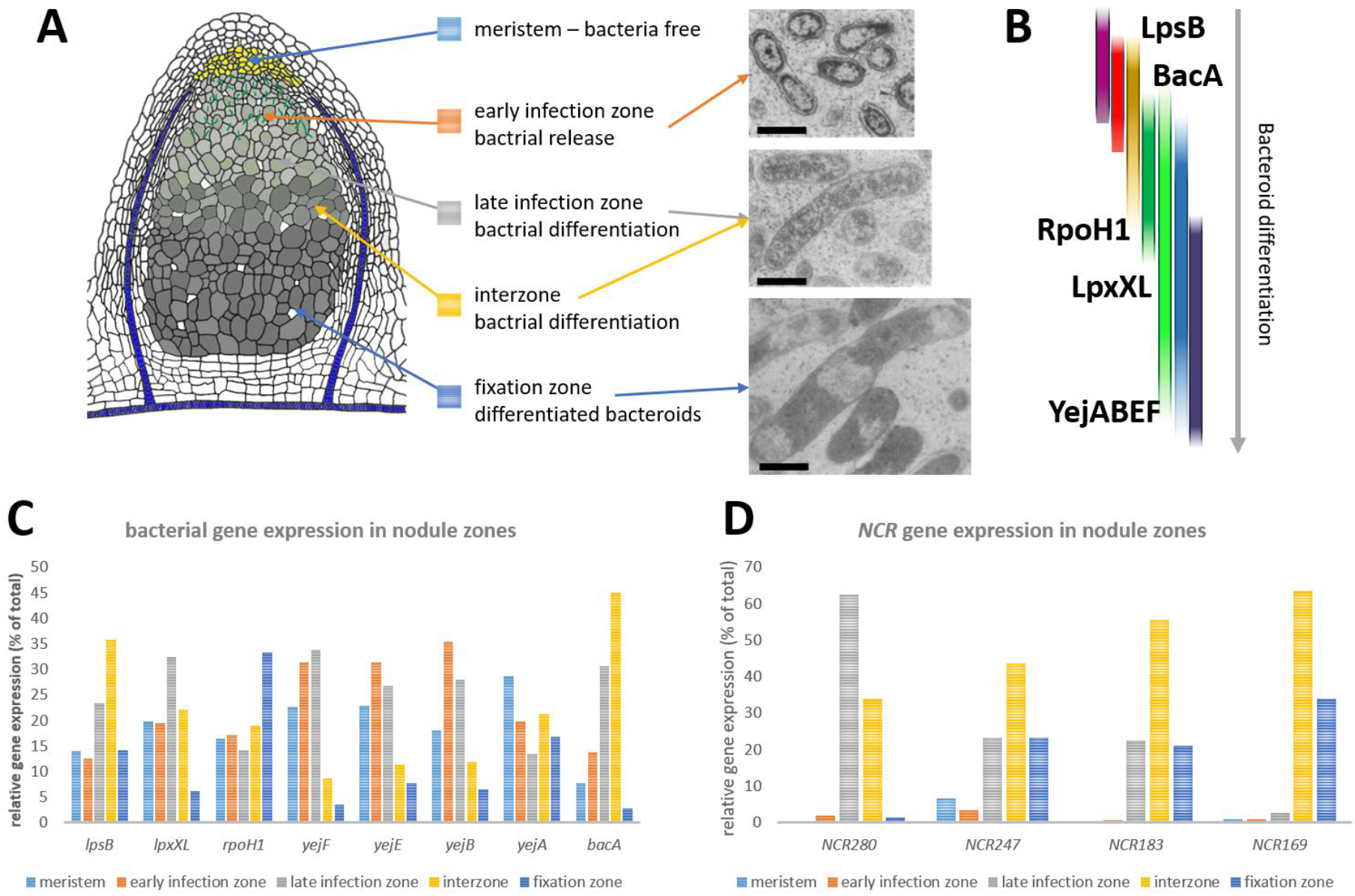
Expression pattern in nodules of bacterial and *NCR* genes. **A**. Schematic drawing of a *Medicago* nodule organized in functional zones. The meristem is bacteria-free and contains dividing cells allowing the organ to grow. In the early infection zone, bacteria proliferate in infection threads (green lines) and are released by endocytosis inside cells derived from the meristem. In the late infection zone and the so-called interzone, the bacteria differentiate into bacteroids. The fixation zone contains the fully differentiated, nitrogen-fixing bacteroids. The pictures show bacteria inside the nodule cells and are presented at the same scale (bar is 1 µm) allowing to appreciate the transformation of the bacteria. **B**. Against the backdrop of a changing landscape of NCR peptides (rainbow colors representing schematically peptides appearing and disappearing at different stages of symbiotic cell differentiation), the bacterial functions described in this study are critical at distinct stages of the bacteroid differentiation process. It should be noted that the functions that are essential in early stages, such as LpsB and BacA, can also be important in later stages of the bacteroid differentiation. However, the phenotypic analysis of the corresponding mutants cannot reveal these putative late roles. **C**,**D**. The relative expression profile (% of total) of the studied bacterial genes *lpsB, lpxXL, rpoH1, yejF, yejE, yejB, yejA* and *bacA* (**C**) and of *NCR280, NCR247, NCR183* and *NCR169* (**D**) in the meristem, early infection zone, late infection zone, interzone and fixation zone are displayed. Data was extracted from (15) and was obtained by RNA-seq analysis on laser-microdissected nodule tissues (33).

**Figure S2.**
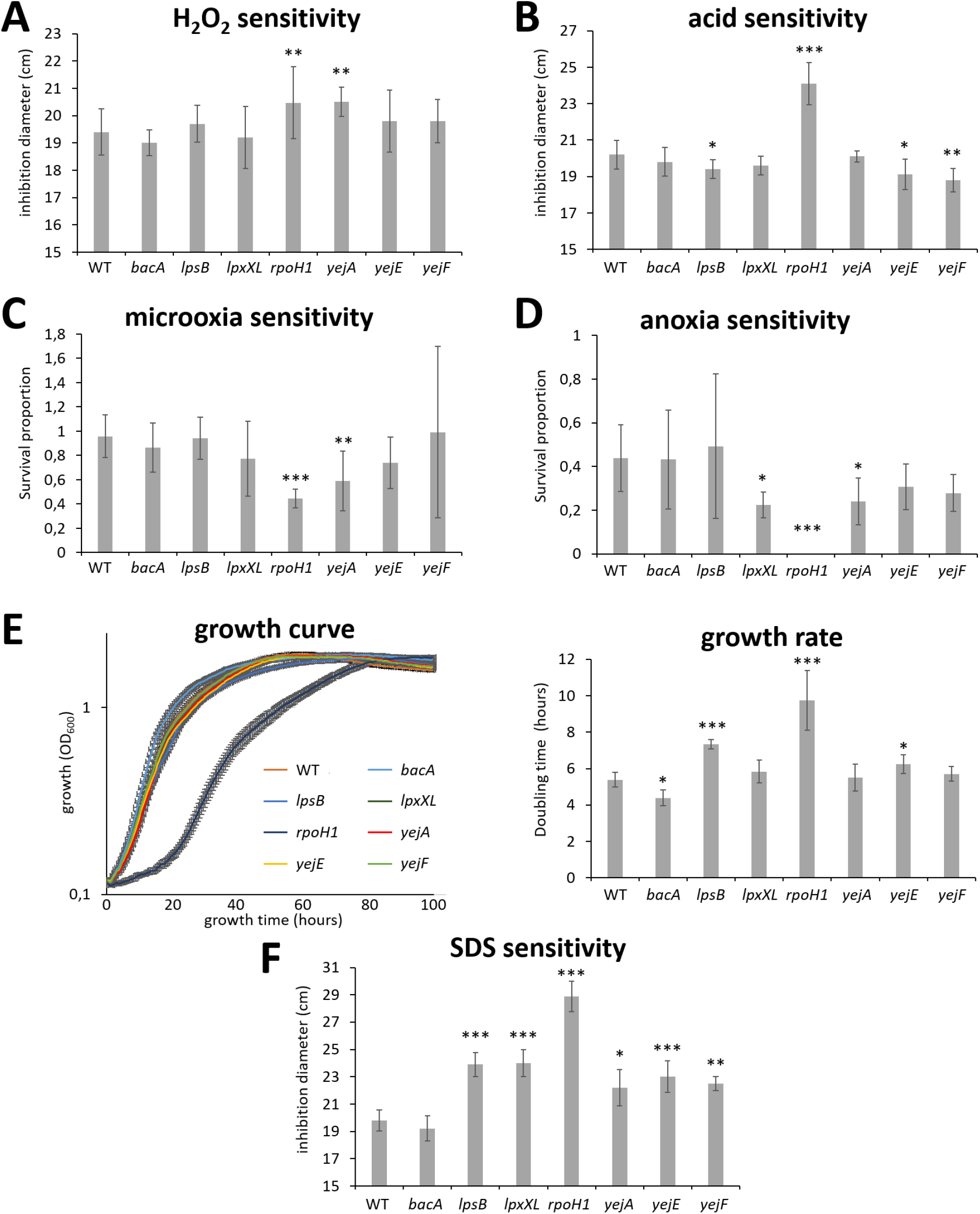
Phenotypes of *Sinorhizobium meliloti* mutants in *in vitro* stress conditions. **A**. Growth inhibition in the presence of H_2_O_2_ (n=10). **B**. Growth inhibition in the presence of HCl (n=10). **C**. Growth inhibition in microoxia (n=4). **D**. Growth inhibition in anoxia (n=4). **E**. Growth curves and growth rate in liquid medium (n=10). **F**. Growth inhibition in the presence of SDS (n=10). Growth inhibition in (**A**,**B**,**F**) was determined by the size of a halo formed around a paper patch soaked in the stress compound and in (**C**,**D**) by colony forming units counting. Error bars in all panels are standard deviations. Nonparametric Kruskal-Wallis and post-hoc Dunn tests were used to assess the significance of differences in the data of panels A to F (* is *p*<0.05; ** is *p*<0.01; *** is *p*<0.001). One representative experiment out of at least two is shown.

**Figure S3.**
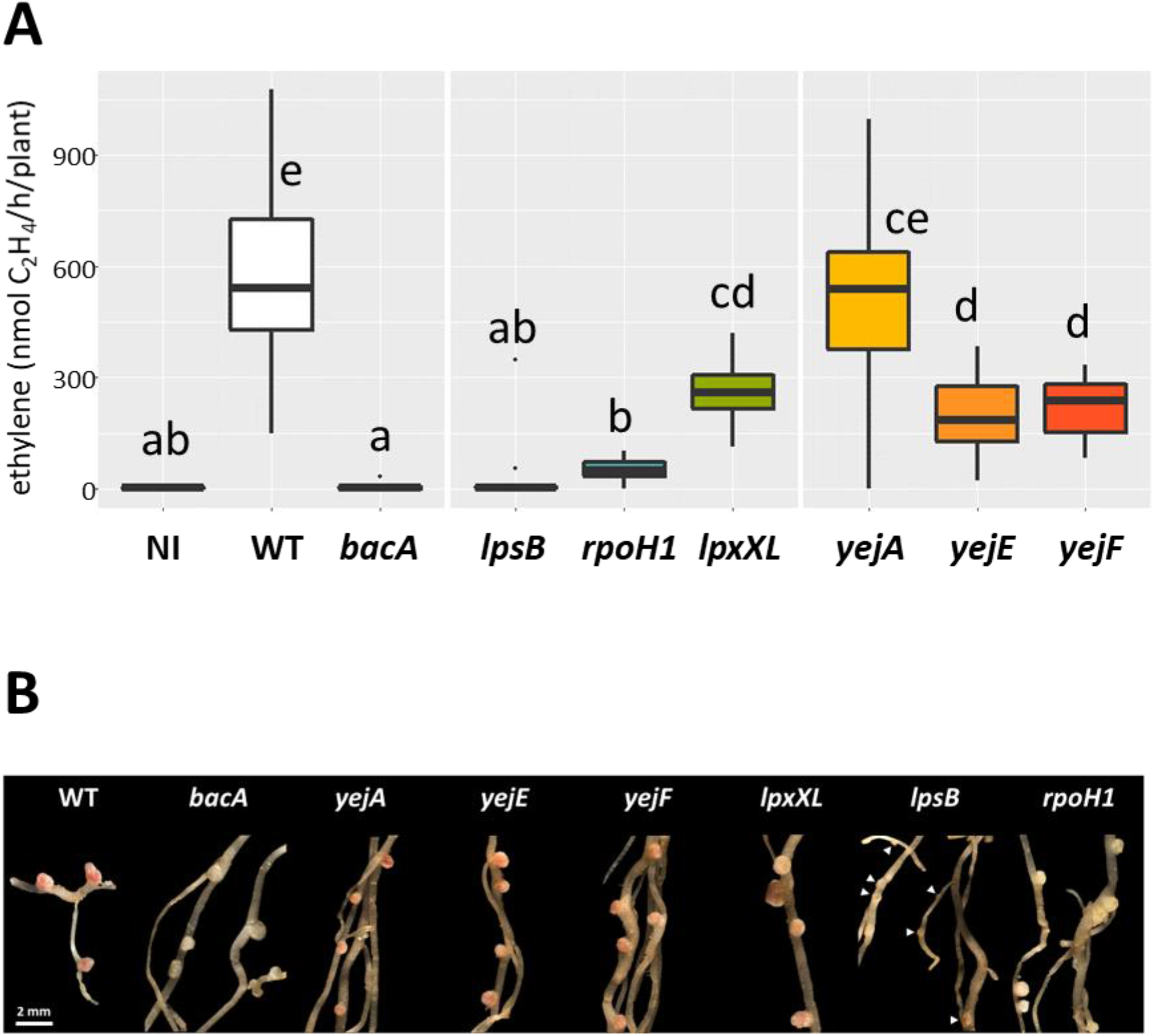
Symbiotic phenotypes of *Sinorhizobium meliloti* NCR-sensitive mutants in *Medicago truncatula*. **A**. Nitrogen fixation activity determined by the acetylene reduction assay on whole roots of nodulated plants infected with the indicated bacterial mutants at 21 days post inoculation. NI, non-inoculated control plants; WT, plants nodulated by the wild-type strain Sm1021. Boxplots were generated from 15 plants each. Letters associated with each condition represent statistically different classes determined by a non-parametric Dunn test, with a α threshold equal to 0.05. **B**. Nodule phenotypes at 21 days post inoculation. Arrowheads indicate small nodules elicited by the *lpsB* mutant. Scale bar (2 mm) applies to all panels.

**Figure S4.**
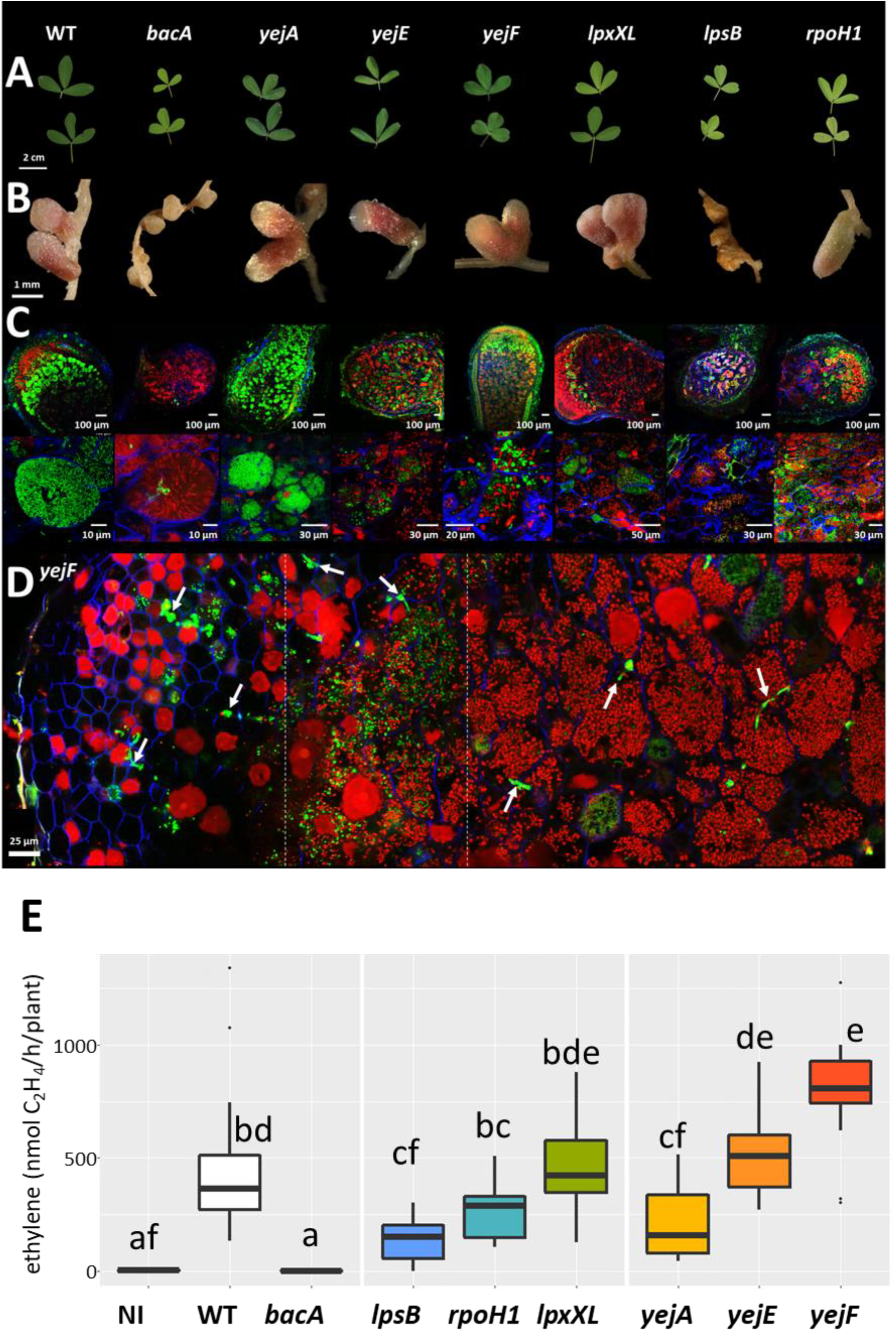
Symbiotic phenotype of *Sinorhizobium meliloti* mutants during symbiosis with *Medicago sativa*. **A**. Leaves of plants inoculated with the indicated bacterial strains. Scale bar = 2 cm. **B**. Nodule phenotype at 21 days post inoculation. Scale bar = 1 mm. **C**. Bacteroid viability determined by live-dead staining of nodule sections and confocal microscopy. Top row images, full nodule sections; Bottom row images, enlarged images of symbiotic cells. Scale bars are indicated in each panel. **D**. Composite image of a *yejF*-infected nodule section showing the rapid permeabilization of the membranes of the internalized bacteria (red staining). Note the bacteria in infection threads (white arrows) are not permeabilized (green staining). The images of the composition are separated by dashed lines. **E**. Nitrogen fixation activity determined by the acetylene reduction assay on whole roots of nodulated plants infected with the indicated bacterial mutants at 21 days post inoculation. NI, non-inoculated control plants; WT, plants nodulated by the wild-type strain Sm1021. Boxplots were generated from 15 plants each. Letters associated with each condition represent statistically different classes determined by a non-parametric Dunn test, with a α threshold equal to 0.05.

**Figure S5.**
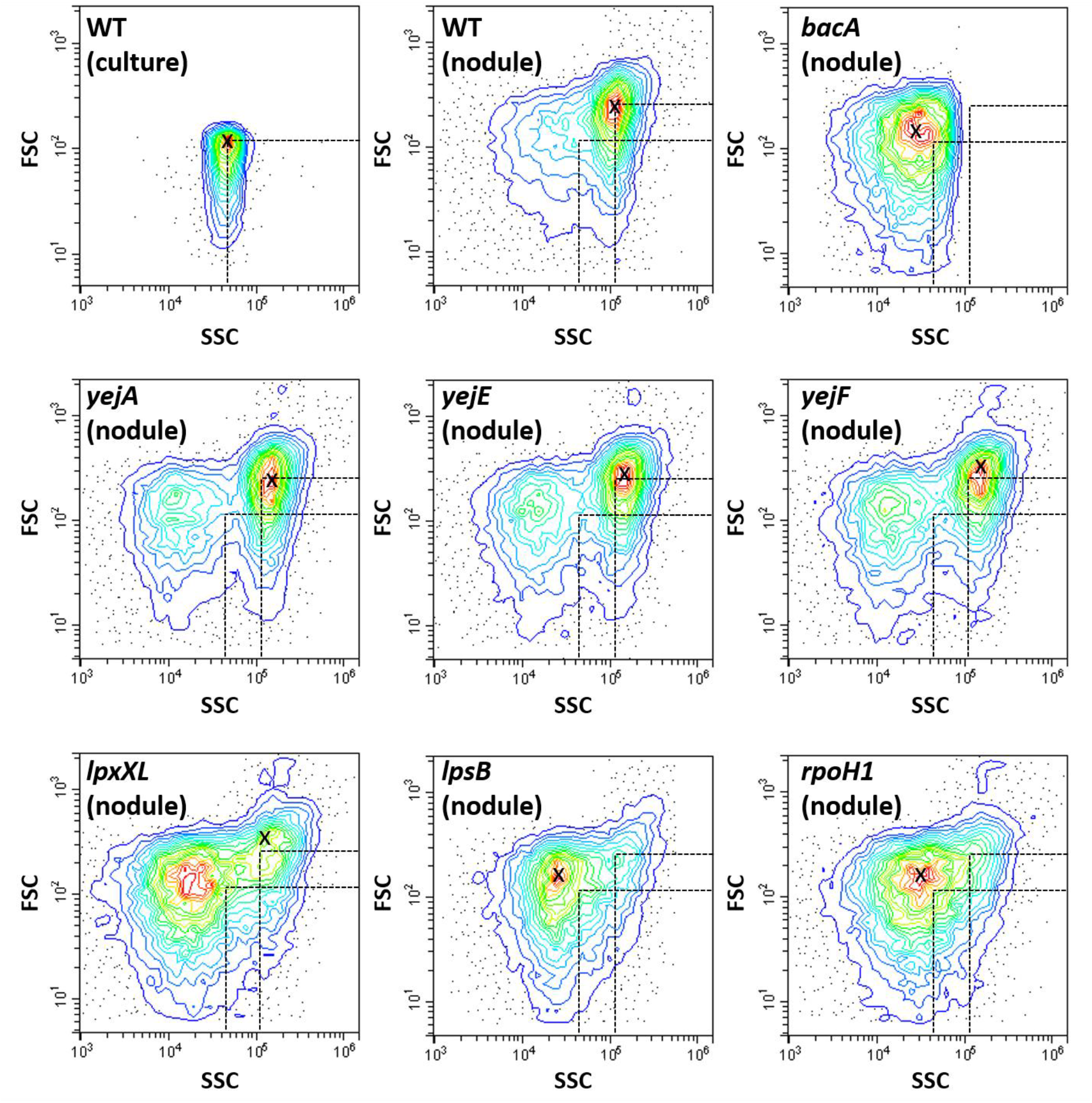
Morphological parameters in nodule bacteria of *Medicago truncatula* nodules. Flow cytometry analysis of the morphology of bacteria isolated from nodules infected with the indicated strains. The forward scatter (FSC, y-axes) is represented in function of the side scatter (SSC, x-axes). The “X” indicates the peak values for each sample. The position of the peak value in the cultured bacteria and in the wild-type bacteroids are indicated in each panel by the hatched lines.

**Figure S6.**
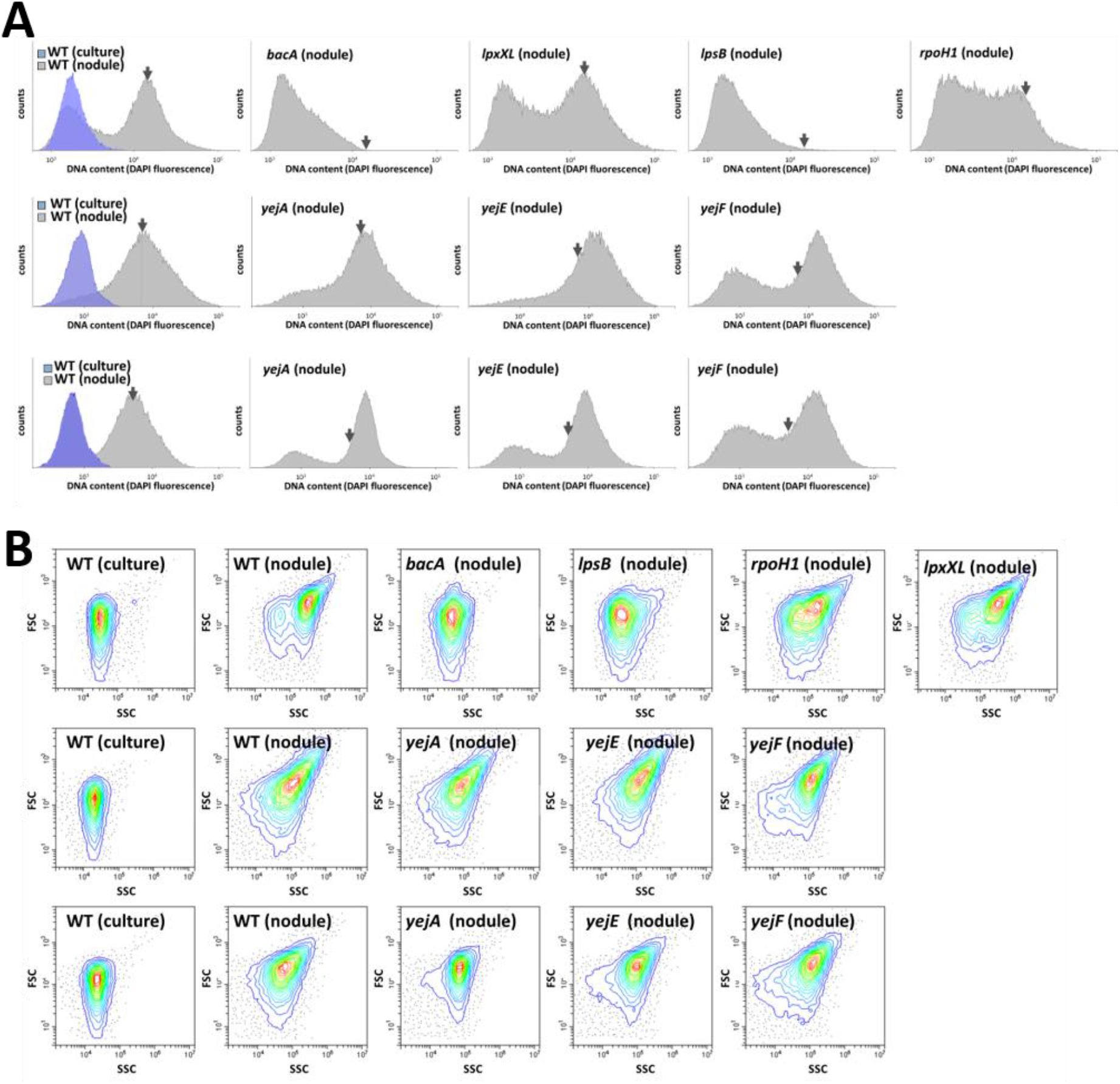
DNA content and morphological parameters in nodule bacteria in *Medicago sativa*. **A**. Flow cytometry analysis of the DNA content of bacteria in culture or isolated from nodules infected with the indicated strains and stained with 4’,6-diamidino-2-phenylindole (DAPI). The cell counts (y-axes) are represented in function of the DAPI fluorescence (x-axes). The arrow in each panel indicates the mean DNA content of wild-type bacteroids. The rows represent three sets of independent experiments and the values of DAPI fluorescence cannot be compared between experiments because of differences in instrument settings. The first two rows are samples prepared from 21 dpi nodules; the third row is from 32 dpi nodules. The latter shows that the phenotype of the *yejA* mutant evolves in older nodules. **B**. Flow cytometry analysis of the morphology of bacteria isolated from nodules infected with the indicated strains. The forward scatter (FSC, y-axes) is represented in function of the side scatter (SSC, x-axes).

**Figure S7.**
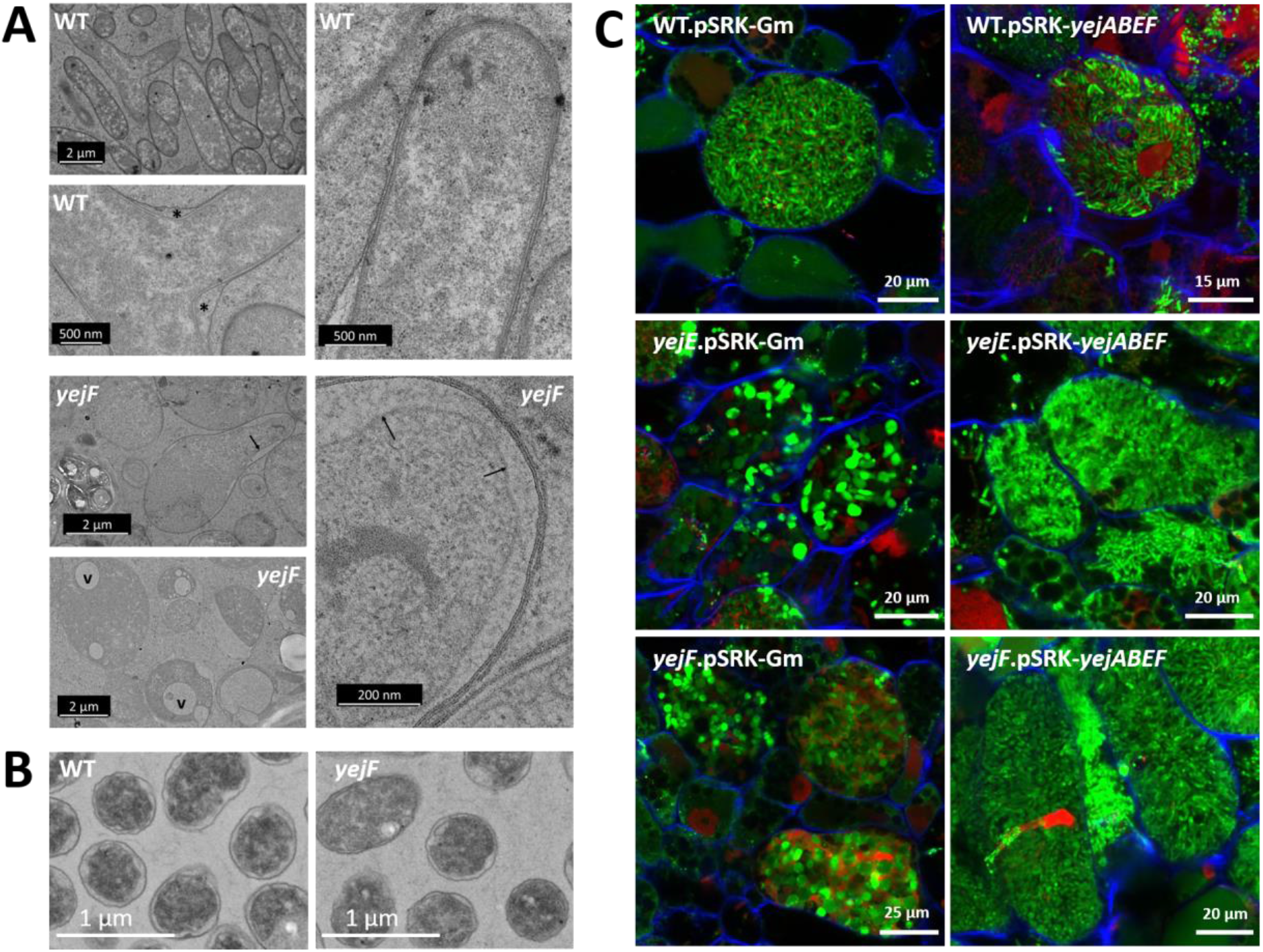
Cellular defects in the *yejF* mutant bacteroids in *Medicago sativa* nodules. **A**. Transmission electron microscopy of wild type (WT) and *yejF* mutant bacteroids. The arrows indicate retracted inner membranes in the *yejF* bacteroids, “v” indicate vacuoles and “*” indicate an enlarged peribacteroid space found in the angle of a branched bacteroid. **B**. Ultrastructure of wild type (WT) and the *yejF* mutant bacteria in culture by transmission electron microscopy. Scale bars are 1 µm. **C**. Confocal microscopy of sections of nodules infected with wild type (WT), *yejE* or *yejF* mutant bacteria carrying the empty plasmid pSRK-Gm or the complementing plasmid pSRK-*yejABEF*.

**Figure S8.**
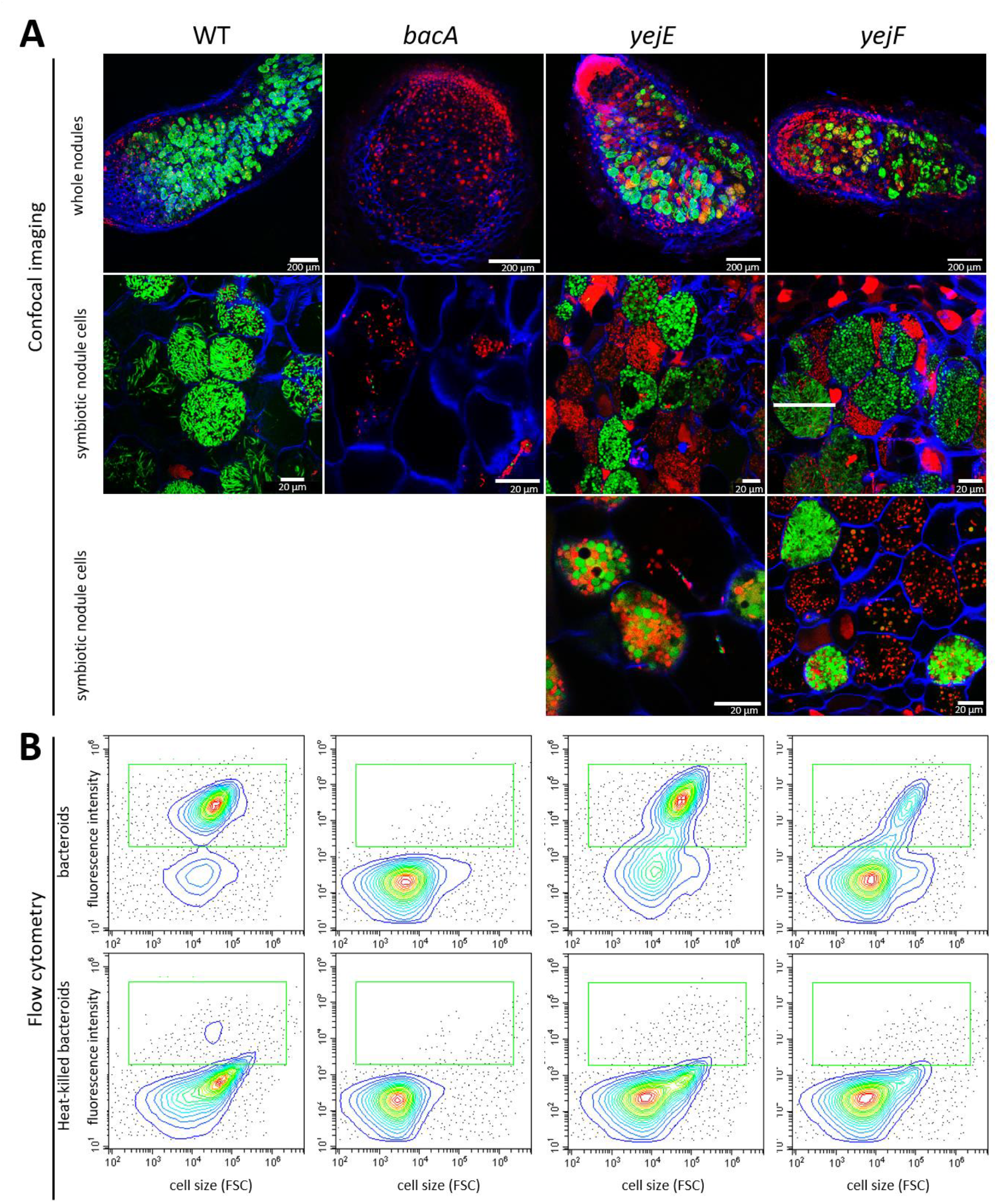
Nitrogenase expression in the *yejE* and *yejF* mutant bacteroids in *Medicago sativa* nodules. **A**. Confocal microscopy of sections of nodules infected with *S. meliloti* Sm1021.p*pnifH*-*GFP, bacA*.p*pnifH*-*GFP, yejE*.p*pnifH*-*GFP* or *yejF*.p*pnifH*-*GFP* and stained with propidium iodide (red stain). Green stained bacteroids are functional while red stained bacteroids are non-functional. **B**. Flow cytometry determination of GFP levels in nodule bacteria (upper panels) and heat-killed nodule bacteria (lower panels). The green square shows the position of the GFP-positive bacteroids. FSC is forward scatter.

**Table S1.** Survival after NCR treatment of wild-type and mutant *Sinorhizobium meliloti*.

| peptide<br>(concentration) | survival (%)<br>relative to<br>untreated<br>bacteria<br>(n=3) | Standard<br>deviation | survival<br>relative to<br>WT (%) | p-value<br>(Dunn test) |
| --- | --- | --- | --- | --- |
| **NCR169 (10 µM)** |  |  |  |  |
| WT | 8,73 | 3,41 | 100,00 |  |
| *bacA* | 4,80 | 4,43 | 55,00 | 0,05 |
| *yejA* | 10,91 | 8,52 | 125,00 | 0,30 |
| *yejE* | 9,45 | 1,70 | 108,33 | 0,26 |
| *yejF* | 13,67 | 8,86 | 156,67 | 0,23 |
| *lpxXL* | 1,02 | 0,34 | 11,67 | 0,02 |
| *lpsB* | 6,55 | 1,70 | 75,00 | 0,48 |
| *rpoH* | 7,56 | 0,68 | 86,67 | 0,23 |
| **NCR183 (10 µM)** |  |  |  |  |
| WT | 0,05 | 0,03 | 100,00 |  |
| *bacA* | 0,01 | 0,01 | 12,50 | 0,24 |
| *yejA* | 0,13 | 0,11 | 275,00 | 0,27 |
| *yejE* | 0,00 | 0,00 | 0,00 | 0,02 |
| *yejF* | 0,00 | 0,00 | 0,00 | 0,02 |
| *lpxXL* | 0,00 | 0,00 | 0,00 | 0,02 |
| *lpsB* | 0,00 | 0,01 | 5,00 | 0,12 |
| *rpoH* | 0,23 | 0,15 | 487,50 | 0,15 |
| **NCR247 (25 µM)** |  |  |  |  |
| WT | 0,03 | 0,02 | 100,00 |  |
| *bacA* | 0,00 | 0,00 | 3,85 | 0,08 |
| *yejA* | 0,05 | 0,00 | 153,85 | 0,37 |
| *yejE* | 0,00 | 0,00 | 3,85 | 0,03 |
| *yejF* | 0,00 | 0,00 | 0,00 | 0,01 |
| *lpxXL* | 0,00 | 0,00 | 0,00 | 0,01 |
| *lpsB* | 0,00 | 0,00 | 0,00 | 0,01 |
| *rpoH* | 0,00 | 0,00 | 0,00 | 0,01 |
| **NCR280 (25 µM)** |  |  |  |  |
| WT | 0,05 | 0,01 | 100,00 |  |
| *bacA* | 0,01 | 0,02 | 23,91 | 0,23 |
| *yejA* | 0,00 | 0,00 | 2,17 | 0,02 |
| *yejE* | 0,00 | 0,00 | 0,00 | 0,01 |
| *yejF* | 0,01 | 0,01 | 10,87 | 0,14 |
| *lpxXL* | 0,00 | 0,00 | 0,00 | 0,01 |
| *lpsB* | 0,00 | 0,00 | 0,00 | 0,01 |
| *rpoH* | 0,07 | 0,02 | 134,78 | 0,29 |
| **NCR247 (25 µM)** |  |  |  |  |
| WT. pSRK-Gm | 0,16 | 0,04 | 100,00 |  |
| *yejF*. pSRK-Gm | 0,00 | 0,00 | 0,12 | 0,01 |
| WT. pSRK-*yejABEF* | 0,14 | 0,04 | 87,99 | 0,36 |
| *yejF*. pSRK-*yejABEF* | 0,19 | 0,08 | 117,20 | 0,36 |

**Table S2.**
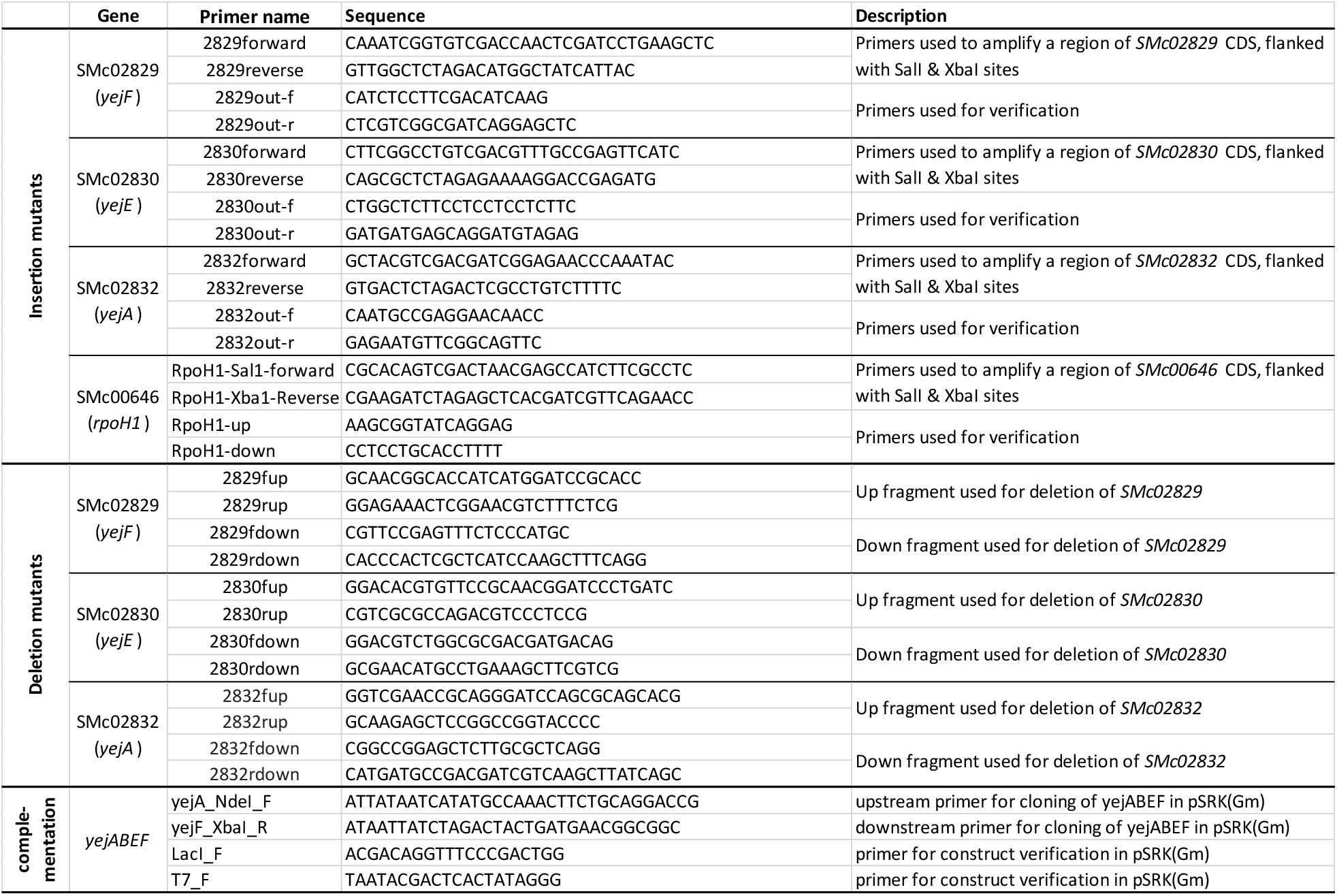
Primers used for mutant constructions.

